# Increased temporal discounting and reduced model-based control in problem gambling are not substantially modulated by exposure to virtual gambling environments

**DOI:** 10.1101/2021.09.16.459889

**Authors:** Luca R. Bruder, Ben Wagner, David Mathar, Jan Peters

**Affiliations:** Department of Psychology, Biological Psychology, University of Cologne, Germany

**Author notes:** **Contact: Luca Bruder**, **Jan Peters**. **Author Contributions** Conceptualization: J. P., L. B. Data curation: L. B. Formal analysis: L. B., B.W., D. M. Funding acquisition: J. P. Investigation: L. B. Methodology: J. P., L. B. Project administration: L. B. Writing – original draft: L. B. Writing – review & editing: J. P., L. B., B. W., D. M. **Additional Information** The authors declare no competing interests.

## Abstract

High-performance virtual reality (VR) technology has opened new possibilities for the examination of the reactivity towards addiction-related cues (cue-reactivity) in addiction. In this preregistered study (https://osf.io/4mrta), we investigated the subjective, physiological, and behavioral effects of gambling-related VR environment exposure in participants reporting frequent or pathological gambling (n=31) as well as non-gambling controls (n=29). On two separate days, participants explored two rich and navigable VR-environments (neutral: café vs. gambling-related: casino/sports-betting facility), while electrodermal activity and heart rate were continuously measured using remote sensors. Within VR, participants performed a temporal discounting task and a sequential decision-making task designed to assess model-based and model-free contributions to behavior. Replicating previous findings, we found strong evidence for increased temporal discounting and reduced model-based control in participants reporting frequent or pathological gambling. Although VR gambling environment exposure increased subjective craving, there was if anything inconclusive evidence for further behavioral or physiological effects. Instead, VR exposure substantially increased physiological arousal (electrodermal activity), across groups and conditions. VR is a promising tool for the investigation of context effects in addiction, but some caution is warranted since effects of real gambling environments might not generally replicate in VR. Future studies should delineate how factors such as cognitive load and ecological validity could be balanced to create a more naturalistic VR experience.

## Introduction

Goal-directed decision-decision making is a key aspect of optimal behavior in a complex environment. Alterations therein in the form of prepotent, impulsive response patterns and habitual tendencies may result in adverse consequences in the long run. A prominent example for this are addiction related disorders such as substance-use-disorders ^[1–3]^ or behavioral addictions like gambling disorder (GD)^[4–6]^. Assessing the processes underlying decision making in general and impaired decision making specifically is difficult, because often not even the agent itself knows why a decision was made. A tool to study the processes that underly decision-making impairments is computational psychiatry^[7]^. The young field of computational psychiatry employs theoretically grounded mathematical models to quantify these processes and assess how they are perturbed in psychiatric disorders^[8]^, with the aim of establishing common (transdiagnostic) computational markers ^[9]^. This might in turn inform the development of effective interventions and/or treatment targets and might support the identification of vulnerable individuals.

Studies investigating maladaptive decision-making have identified several related but distinct processes that play a role across psychiatric disorders. One process is the discounting of reward value over time (temporal discounting), as both steep and shallow discounting are associated with different psychiatric conditions^[1]^. In temporal discounting tasks, participants repeatedly choose between a fixed immediate reward and larger rewards that are temporally delayed^[10]^. The degree of temporal discounting is then estimated from choices and/or response time (RT) distributions via computational models. Altered temporal discounting is suspected to be a transdiagnostic marker for several psychiatric disorders^[1,9]^, with addictions and related disorders forming a prominent example^[9,11]^.

Another cognitive process that has received considerable attention for quantifying goal-directed control of decision-making is reinforcement learning (RL)^[12]^. RL is thought to depend on two systems, a habitual “model-free” system that learns stimulus-response associations, and a goal-directed “model-based” system that computes action consequences via a model of the environment^[13]^. The degree to which participants use goal-directed or habitual RL is often assessed with the 2-step task^[14,15]^, a two-stage decision-making task where first stage decisions probabilistically determine the presented second stage, and thereby the rewards that can be obtained. Reduced model-based RL is associated with a range of subclinical symptoms^[16]^ and addiction related disorders including GD^[17]^ and substance use disorder^[18]^.

Prominent characteristics of addiction are compulsive drug seeking and insensitivity to negative consequences^[19]^. Incentive-sensitization theory^[20–22]^ postulates that neural circuits mediating the incentive motivation to obtain rewards become sensitized to reward-predictive cues, giving rise to craving and drug-seeking behavior. These effects are thought to be mediated by the mesocorticolimbic dopamine system^[22]^. For example, exposure to addiction-related cues is correlated with the modulation of striatal value signals during temporal discounting^[6]^, and increases striatal dopamine release in humans^[23]^. Across substance-use disorders and behavioral addictions, such effects are referred to as *cue-reactivity*^[24–26]^. Cue-reactivity manifests on a physiological and subjective level^[24,25]^. Additionally, exposure to addiction-related cues might increase temporal discounting^[6,27,28]^, modulate risk-taking^[29]^ and impair cognitive performance^[30]^.

Cue-reactivity in GD has been examined using visual cues^[4,6,29,31–38]^ or real-life gambling environment exposure^[27,28]^. Both methods arguably represent extremes on the spectrum between highly controlled laboratory environments and field studies. Field studies have high ecological validity but lack control over confounding factors and complicate the assessment of physiological variables. Conversely, lab studies yield control over confounding variables but lack ecological validity. We recently proposed a virtual reality (VR) approach that combines ecological validity with the advantages of a highly controlled lab environment^[39–41]^. VR allows the concurrent measurement of physiological, subjective, and behavioral cue-reactivity in an ecologically valid virtual environment. Participants are equipped with head-mounted displays and immersed in two rich and navigable VR environments: a (neutral) café or a (gambling-related) casino. Participants explore the environments and subsequently perform behavioral tasks within them.

We have shown that behavioral data obtained in VR yield reliable estimates of temporal discounting^[39]^, and allow for a comprehensive RT-based modeling via the drift-diffusion model (DDM)^[39,42,43]^. Here, the decision process is modelled as a dynamic diffusion process between two boundaries, providing both a more detailed account of the underlying latent decision processes^[44–48]^ and more stable parameter estimates^[49,50]^. Recent studies have successfully applied this approach to disentangle dopamine effects on temporal discounting^[48]^, quantify reinforcement learning impairments in GD^[51]^ and clarify effects of medial orbitofrontal cortex lesions on decision-making^[46]^.

This pre-registered VR study had three aims. First, we extended previous VR work on cue-reactivity in addiction ^[40,52,53]^ by comprehensively modeling the effects of gambling-related environments in GD on cognitive processes during decision-making. We hypothesized that participants reporting frequent and/or pathological gambling behavior would show overall increased temporal discounting^[9,11]^ and reduced model-based RL^[17]^ compared to non-gambling controls. Additionally, we predicted that exposure to a VR gambling context would increase subjective craving (urge-to-gamble), further increase temporal discounting^[5,10]^ and further reduce model-based RL^[17]^ in GD compared to controls. We hypothesized that physiological cue-reactivity in GD would manifest as increased heart rate and skin-conductance responses during exposure to a VR gambling context.

## Methods

### Participants

Thirty-one participants (three female) reporting regular gambling and at least one DSM-V^[19]^ criterion for gambling disorder aged between 20 and 41 (mean = 26.06, std = 5.43) as well as thirty non-gambling control participants (three female) in a matched age range (mean = 26.83, std = 4.65) were invited to the lab on three different testing days. Groups were additionally matched on years of education and smoking (Table 1). Participants were recruited via flyers posted at local gambling venues and via postings in local internet forums. No participant reported a history of traumatic brain injury, psychiatric or neurological disorders or severe motion sickness.

**Table 1.** Summary of group characteristics. p-values printed in bold font are significant at .05.

| Screening | Gambling Group | Control Group | Group Difference |
| --- | --- | --- | --- |
| Gender (male/ female) | 28/3 | 26/3 | $X^2(1) < 0.001$,<br>$p = 1$ |
| Age | 26.06 (5.34) | 26.83 (4.65) | $U = 426$,<br>$p = 0.73$ |
| Years of education | 12.1 (1.68) | 12.07 (1.25) | $U = 454.5$,<br>$p = 0.94$ |
| DSM-V criteria for GD<sup>[19]</sup> | 5.32 (2.41) | 0 (0) | $U = 899$,<br>**$p < 0.001$** |
| Beck Depression Inventory (BDI)<sup>[54]</sup> | 13.45 (10.08) | 6.45 (5.53) | $U = 639$,<br>**$p < 0.01$** |
| Symptom-Checklist 90 (SCL90)<sup>[55]</sup> | 0.93 (0.7) | 0.34 (0.29) | $U = 684$,<br>**$p < 0.001$** |
| South Oaks Gambling Screen (SOGS)<sup>[56]</sup> | 8.23 (4.55) | 0.24 (0.83) | $U = 873.5$,<br>**$p < 0.001$** |
| Kurzfragebogen zum Glücksspielverhalten (KFG)<sup>[57]</sup> | 24.55 (13.61) | 3.21 (11.3) | $U = 851$,<br>**$p < 0.001$** |
| Alcohol Use Disorder Identification Test (AUDIT)<sup>[58]</sup> | 9.84 (8.87) | 4.79 (3.08) | $U = 580.5$,<br>$p = 0.052$ |
| Fagerström Test for Nicotine Dependence (FTND)<sup>[59]</sup> | 1.32 (2.5) | 0.69 (1.37) | $U = 483$,<br>$p = 0.52$ |
| Gambling-Related Cognitions Scale (GRCS)<sup>[60]</sup> | 70.35 (26.25) | 26.55 (8.14) | $U = 878$,<br>**$p < 0.001$** |
| Gamblers' Beliefs Questionnaire (GBQ)<sup>[61]</sup> | 30.29 (19.32) | 26.21 (15.89) | $U = 483$,<br>$p = 0.62$ |

Participants provided informed written consent prior to their participation, and the study procedure was approved by the Ethics Board of the Germany Psychological Society. The procedure was in accordance with the 1964 Helsinki declaration and its later amendments.

### VR-Setup

The VR-environments were presented using a wireless HTC VIVE head-mounted display (HMD). The setup provided a 110° field of view, a 90 Hz refresh rate, and a resolution of 1440 × 1600 Pixel per eye. Participants had an area of about 6m^2^ open space to navigate the virtual environment. For the execution of the behavioral tasks and additional movement control participants held one VR-controller in their dominant hand. The VR-software was run on a PC with the following specifications: CPU: Intel Core i7-3600, Memory: 32.0 GB RAM, Windows 10, GPU: NVIDIA GeForce GTX 1080 (Ti). The VR-environments themselves were designed in Unity. Auditory stimuli were presented using on-ear headphones.

### VR-Environments

The VR set-up consisted of two environments, one neutral environment (VR_neutral_) and one gambling-related environment (VR_gambling_). Environments contained an identical starting area and different experimental areas. Participants were placed in the middle of a small rural shopping street with a small park adjacent to it. Participants heard low street noises. The starting area was intended to familiarize participants with VR and control for possible confounding effects of entering VR. From this starting area participants could move to the experimental area by entering one of the houses on the shopping street. The entrances were in the same location for both environments. The actual experimental area differed for the two VR environments (see Figure 1). In the VR_neutral_ environment the experimental area comprised a small café including customers and buffet (Figure 1b, c). Participants were surrounded by low conversation and music. In the VR_gambling_ environment the experimental participants were presented a small casino containing slot machines and a sports betting room (Figure 1e, f). Participants could hear slot machine sounds and sports. The floorplan of the VR_gambling_ experimental area was a mirrored version of the VR_neutral_ experimental areas floorplan (Figure 1a, d). Both environments contained eight animated human avatars that performed non-repetitive movements like gambling or ordering food.

**Figure 1.**
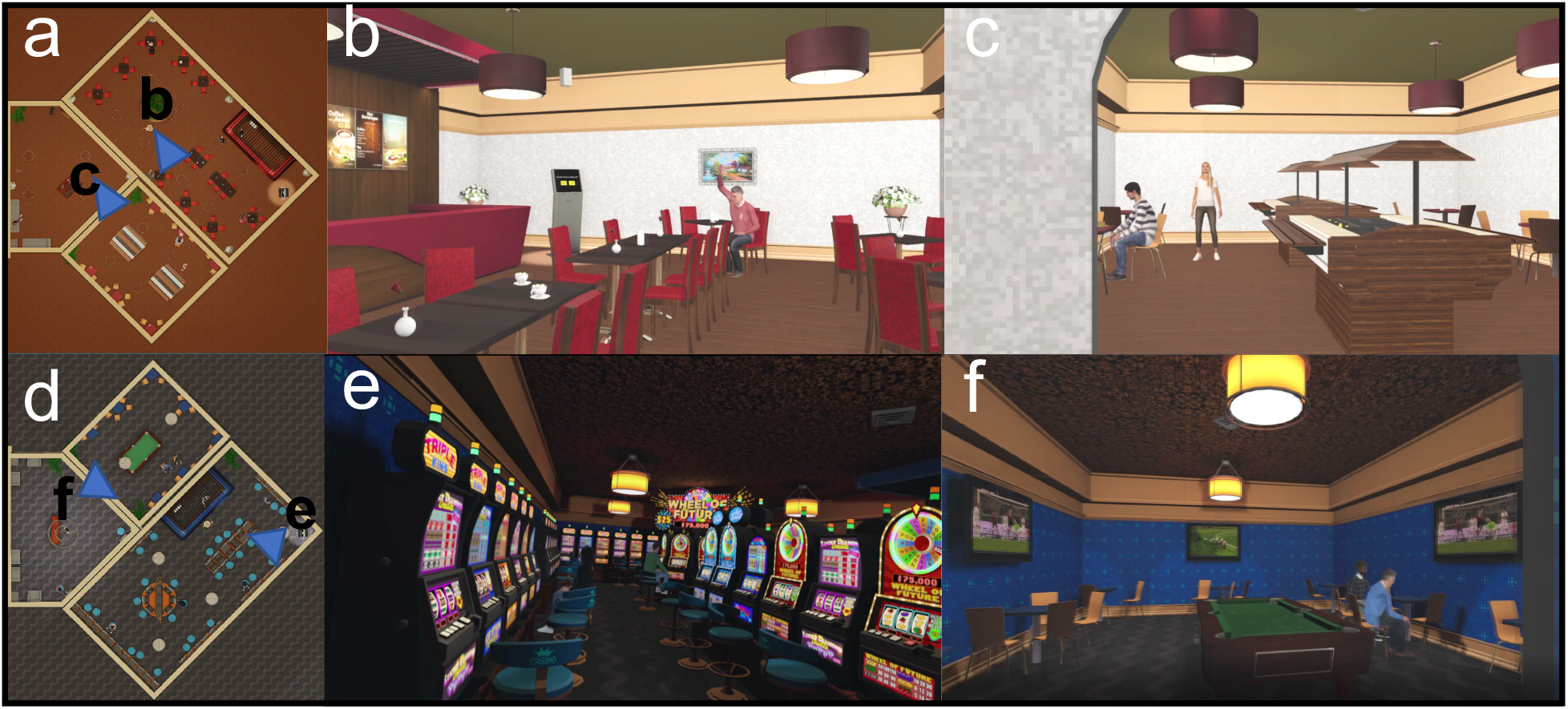
Experimental areas of the VR-environments a) Floorplan of the café within the VR-neutral environment b) View of the main room of the café c) View of the buffet area of the café d) Floorplan of the casino within the VR-gambling environment e) View of the main room of the casino f) View of the sports bar within the casino

### Experimental procedure

Participants were invited for three testing sessions on three different days. The mean interval between sessions was 2.06 days with a range from 1 to 7 days. During the first session, participants complete a range of questionnaires and working memory tasks (see baseline screening). This session took approximately two and a half hours. In the second and third session participants entered one of the two VR environments. The order of the two VR environments was counter-balanced across participants. Upon arrival at the lab for the VR-sessions participants were first introduced to the VR equipment and handling. Subsequently they received detailed instructions for the behavioral tasks to be performed in VR. Participants were then seated, and a five-minute baseline measurement of physiological measures was obtained (baseline phase). The experimenter then helped the participants to apply the VR headset. Upon VR immersion, participants found themselves in the starting (outdoor) area of the VR environment and were instructed to explore it for five minutes (first exploration phase). Participants were then instructed to enter the interior experimental area of the VR environment and explore it for an additional five-minute period (second exploration phase). Experimental phases were each divided into five one-minute bins (B1 to B5 for the baseline phase, F1 to F5 for the first exploration phase and S1 to S5 for the second exploration phase). Following S5, participants were asked to proceed to the behavioral tasks, which were presented on a terminal within the experimental area.

### Physiological measurements

Electrodermal activity (EDA)^[62]^ was measured using a BioNomadix-PPGED wireless remote sensor together with a Biopac MP160 data acquisition system (Biopac Systems, Santa Barbara, CA, USA). A GSR100C amplifier module with a gain of 5V, low pass filter of 10 Hz and a high pass filter DC were included in the recording system. The system was connected to the acquisition computer running under Biopac’s AcqKnowledge software. Triggers for the events within the VR-environments were send to the acquisition PC via digital channels from the VR-PC. Disposable Ag/AgCl electrodes were attached to the thenar and hypothenar eminences of the non-dominant palm. Isotonic paste (Biopac Gel 101) was used to ensure optimal signal transmission. The signal was measured in micro-Siemens units (mS). The same Biopac MP160 system with a BioNomadix-PPGED wireless remote sensor was used to record the heart rate at the fingertip.

### Temporal discounting task

Participants completed two behavioral tasks in each VR environment: a temporal discounting task^[10]^ and the Two-Step task^[14,15]^ (a sequential RL task). The temporal discounting task consisted out of 96 choices between an immediate (smaller-but-sooner, SS) reward fixed at 20 Euros, and larger but delayed (larger-later, LL) rewards. LL options were created by multiplying the SS option with a range of factors (range 1.025 to 3.85) combined with different temporal delays (range 1 to 122 days). Two sets of LL options were created by taking all possible combinations of six delays and 16 factors for each set. The two sets were matched for mean LL amount and mean delay. The order of presentation of the two sets was counterbalanced across participants. Participants were informed that one trial per session would be randomly selected and paid out in form of voucher for a widely known online store.

Options per trial were presented in two yellow squares on a black background on a display positioned within the experimental area of the VR environments (Figure 2). Offers were randomly assigned to the right or left side of the virtual display. Participants could take as long as they wanted to make a response, but they were instructed to decide intuitively. After a decision was made by aiming at the preferred option with the VR controller and pulling the trigger, a short inter-trial-interval (ISI) of .5 to 1 seconds followed. During this ISI the two yellow squares were filled with questions marks. Subsequently the next trial started.

**Figure 2.**
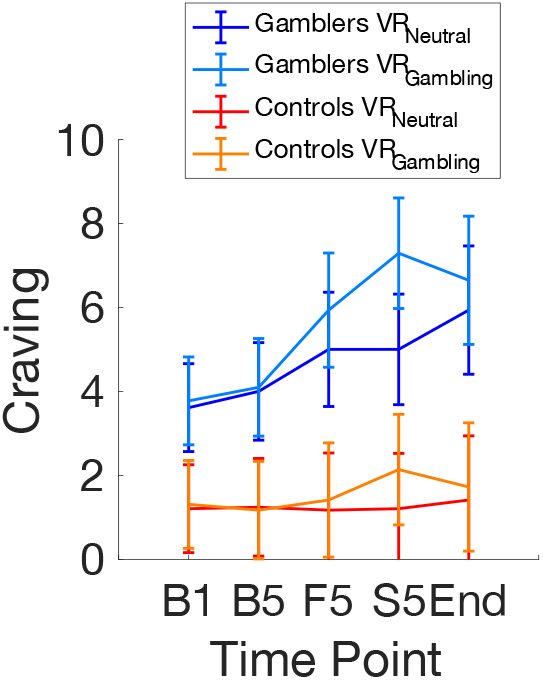
Subjective urge-to-gamble (craving) was verbally reported via a 10-point-likert scale at five time points, at the beginning and end of the pre-VR baseline phase (B1, B5), at the end of the first VR exploration phase (F5, VR outdoor area) and at the end of the second VR exploration phase (S5, VR_neutral_ vs. VR_gambling_) and upon completion of the experimental tasks (End).

### Sequential RL task

Next, participants completed 200 trials of a modified version of the 2-step task proposed by Daw and colleagues^[14]^. The task consists of two stages. On each stage, participants choose between two abstract visual cues. The first stage (S1) consisted of two stimuli, one of which had to be chosen. Depending on the choice in S1, participants were then taken to one of two second stages (S2) that were characterized by different colors and different available stimuli. Here, one of the two S2 stimuli was selected, and rewarded with points ranging from 0 to 99. One S1 cue led to one second stage with a probability of 70% (“common” transition) and to the other with a probability of 30% (“rare” transition). Transitions were reversed for the other S1 option and fixed across trials. Following selection of an S2 option, the obtained points were shown below the chosen picture for one second. S2 rewards were determined by four independent Gaussian random walks with reflecting boundaries (min: 0, max: 99). These walks were pre-computed and counterbalanced across sessions. Participants were informed about the task structure and their comprehension of the task was repeatedly questioned by the experimenter. Additionally, participants completed five example trials to get used to the task timing. In both stages, participants had three seconds to log their response. If they failed to do so they received zero points and the next trial started. Motivation to perform the task correctly was ensured by linking the final score to an additional financial reward that participants could receive. This was done by multiplying the final score of the participant with a factor of .0005 resulting in an additional reward of between five and nine €.

### Baseline screening

During the first session, participants answered a range of questionnaires, and their working memory capacity was probed. A list of the questionnaires can be found in Table 1. To probe the working memory capacity of the participants we applied four established tests. First, participants completed the Rotation Span Task, which tests the ability to memorize a sequence of arrow orientations while being distracted by a letter rotation task^[63]^. Second, participants had to memorize sequences of letters, while doing simple calculations. This Operation Span Task was adapted from the Complex Span Task^[64]^. Third, we tested the listening span of participants by having them remember the last words of sentences presented in variably sized blocks. This task was adapted from the German version of Reading Span Task^[65]^. Fourth, participants performed a forward and a backward version of the Digit Span Task. Participants heard sequences of digits and had to remember them in the same or reversed order^[66]^.

### Temporal discounting: model-agnostic analysis

Temporal discounting data were first analyzed with a model-free approach, without a-priori model assumptions. Points of subjective equivalence between SS and LL rewards (indifference points) were estimated by fitting logistic functions to the choices for each delay. Indifference points were then plotted per participant and the area under the resulting curve (AUC) was calculated following standard procedures^[67]^. AUC values were compared across groups and sessions using a mixed ANOVA.

### Temporal discounting: computational modeling

The effect a delay has on the subjective valuation of a reward can be accurately modelled by a hyperbolic function^[68,69]^. We therefore employed hierarchical Bayesian modeling to determine each participant’s rate of discounting for the different sessions^[8]^. A hierarchical hyperbolic discounting model was fit to data of each group individually (Eq. 1):

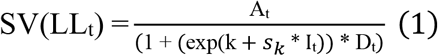

Here, SV(LL_t_) is the subjective (discounted) value of the LL on trial *t*. A and D represent the objective reward amount and the delay to the LL, respectively. The steepness of the hyperbolic discounting function is governed by the parameter *k* (discount rate). To avoid numerical instability of *k*-values close to, we estimated the parameter in log-space (log[k]). The parameter *s*_*k*_ represents the change in *log(k)* from the VR_neutral_ to the VR_gambling_ session. Finally, *I*_*t*_ is a dummy variable coding the experimental session for trial *t*.

The hyperbolic model was subsequently combined with two different choice rules, a softmax action selection rule^[12]^ and the drift diffusion model (DDM)^[45]^. Softmax determines the probability of choosing the LL option on a given trial *t* (Eq. 2):

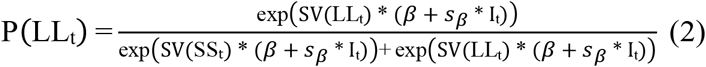

Here, the *β* -parameter determines choice stochasticity with respect to model-based subjective values. For *β* = 0 choices are random, whereas higher *β* values reflect stronger dependency of choices on modeled values. Again, we included a term (*s*_*β*_) modeling condition-dependent changes in decision noise.

The second choice rule we applied was the DDM. In addition to the choices, the DDM includes response times (RTs) in the analysis to decompose RT distributions into latent decision processes. Recently, the DDM has increasingly been applied in the context value-based decision-making, including temporal discounting^[39,46,48]^ and reinforcement learning^[44,45]^ and models two-alternative forced-choice decisions as a noisy evidence accumulation process between two boundaries. As soon as the evidence in favor of one option exceeds the corresponding boundary, the process terminates, and the choice is executed. The upper boundary coded LL choices, whereas the lower boundary coded SS choices. RTs of SS choices were multiplied with -1 prior to the model estimation. We excluded, the 2.5% slowest and fastest trials of each participant from the analysis to prevent outlier trials from negatively impacting model fit^[45,46]^.

Within the DMM, the RT on trial *t* is then distributed according to the Wiener first passage time (*wfpt*) (Eq. 3):

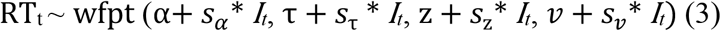

Here *α* represents the boundary separation reflecting the speed-accuracy tradeoff. The non-decision time τ models RT components unrelated to the evidence accumulation process (e.g. perception and motor preparation). The starting point of the diffusion process is modeled via the bias *z*. Finally, the rate of evidence accumulation is described by the drift-rate v. The s-parameters (*s*_α_, *s*_τ_, *s*_*z*_, *s*_*ν*_) again model condition effects on the corresponding parameters, and *I*_*t*_ is the dummy-coded condition predictor for trial *t*.

We compared three different versions of the DDM. First, we fit a null model (DDM_0_) with a constant drift rate^[46,48]^. We then compared two DDMs that incorporated hyperbolic discounting, either via a linear (DDM_L_) or a non-linear (DDM_S_) mapping of trial-wise value differences on drift rates. For the DDM_L_, v_coeff_ maps the value differences onto *v*:

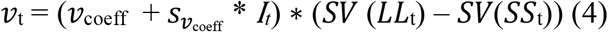

One problem with such a linear mapping is that *v* might increase infinitely with high value differences, leading to a substantial under-prediction of RTs for high value differences^[46]^. As in previous work^[46,48]^, we therefore also examined a non-linear model (DDM_S_) (Eq. 5 and 6)^[44]^:

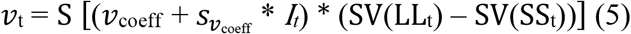

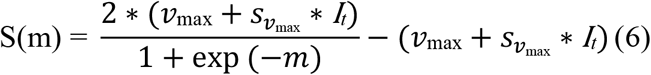

This formulation caps the drift rate at *ν*_max_

We and others have shown that both group- and subject-level DDM_S_ parameters recover well when combined with value-based decision models including temporal discounting^[44,46,48]^.

### 2-step task: model-agnostic analysis

Here we modelled second stage RTs using a hierarchical generalized linear model (HGLM) with the factors transition (common or rare), session (VR_gambling_ or VR_neutral_) and group (gambling or non-gambling control)^[70]^. The 2.5% slowest and fastest trials per each participant were excluded from the analysis to reduce the negative impact of outlier trials on model fit^[45,46]^. We also performed a standard model-agnostic analysis of the 2-step task, modelling the probability to repeat previous trial first stage choices (pStay) as a function of transition, group and previous reward^[14]^. Since the present 2-step task version employed continuous rewards for S2, this analysis was conducted using a moving average of recent rewards to categorize trials into rewarded and unrewarded trials. In addition, we assessed the final score participants achieved in both conditions with a mixed model ANOVA including group and session as factors.

### 2-step task: computational modeling

We again combined an RL model that learns state-action values (Q-values) for options in the different stages with both a softmax and a DDM choice rule. The softmax model was based on the hybrid model proposed by Otto and colleagues^[71]^. Here, both model-free (MF) and model-based (MB) Q-values are tracked for S1 options, whereas S2 options only have MF Q-values. Q_MF_-values for both stages are updated on trial *t* via the prediction error *δ* (Eq. 7 to 10). The subscript j (j ϵ [1,2]) denotes the two possible actions in each stage, while *i* denotes the stage presented.

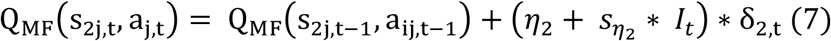

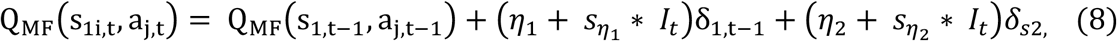

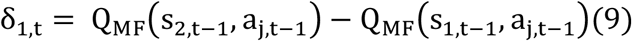

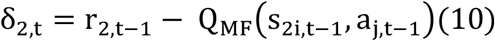

Here, *r* corresponds to the S2 reward obtained on the previous trial. As there are only S2 rewards, S1 Q_MF_-values are based on the Q_MF_-values of the S2 options (equation 11) and on the S2 prediction error. *η*_1_ and *η*_2_ represent the S1 and S2 learning rates (i.e., the impact of prediction errors on future reward expectation). Learning rates were modeled in standard normal space [-4, 4], and back-transformed to the interval [0, 1] via the inverse cumulative normal distribution function.

In contrast, model-based Q-values (Q_MB_, see Eq. 11) take the S_1_-S_2_ transition probabilities as well as S_2_ Q_MF_-values into account:

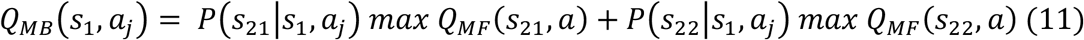

Since there are no further stages to the task, for S2, Q_MF_ = Q_MB_. The S2 softmax choice rule is analogous to the one described for temporal discounting. Here, *β*_2_ governs the choice stochasticity (Eq. 12):

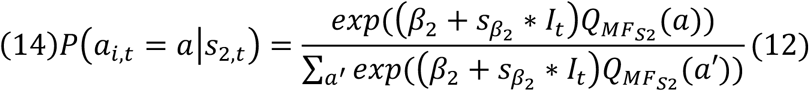

The S1 softmax choice rule then includes separate weighting parameters for Q_MF_ and Q_MB_ (*β*_*MF*_ and *β*_*MB*_) that model the impact of these Q-values on S1 choices (Eq. 13):

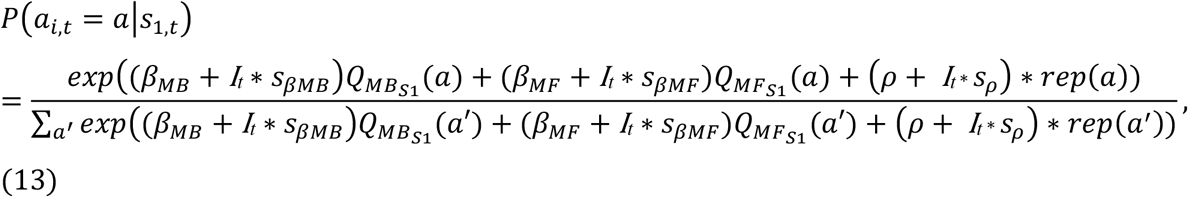

Action selection in S1 is thus modelled by weighing the influence of MB and MF Q-values in a softmax choice rule. Additionally, the parameter *ρ* models perseveration, i.e., the tendency to repeat the previous action. *Rep* takes a value of 1 if the corresponding action was taken on trial t-1, and 0 otherwise. *I*_*t*_ again represents the dummy-coded session.

Finally, we included a term that decayed the Q-values of unchosen stimuli in both stages^[72,73]^ with a decay-rate η_decay_ towards the center of the reward walks (.5) (Eq. 14). For the modelling process the reward values were multiplied by 100 as these values were presented to the participants.

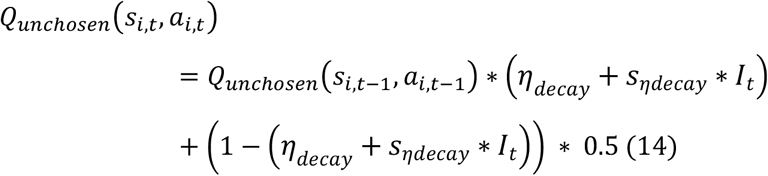

We again extended this modeling approach using the same three DDM choices rules previously described for the temporal discounting task, with the key difference that now trial-wise drift-rates depended on the differences in Q-values between options 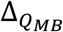 and 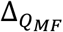 respectively). For S2, this depended only on Q_MF_ value differences (Eq. 15):

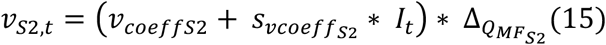

For S1, to take both Q_MF_ and Q_MB_ values into account, we included separate drift-rate coefficients *ν*_coeffMB_ and *ν*_coeffMF_ (Eq. 16):

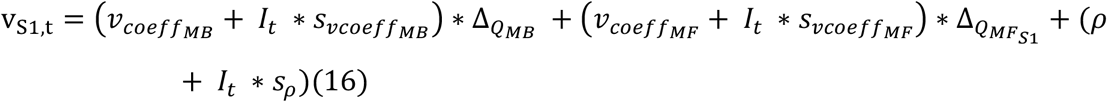

As for the temporal discounting models, the DDM_L_ simply used the drift rates from equations 15 and 16. In contrast, for the DDM_S_ these drift-rates were additionally passed through a sigmoid function S (see Eq. 17):

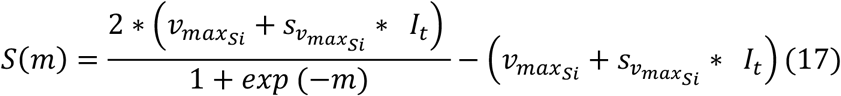

For all models, separate DDM parameters were estimated for the first and second stage of the task.

### Hierarchical Bayesian Models

All models were fit to the data of all participants in a hierarchical Bayesian estimation scheme. Models not including session effects were fit separately per session, yielding separate estimates per participant and session. For group-level means we employed weakly informative uniform or normal priors over numerically plausible ranges (see Supplementary Table 1). For parameters modelling condition effects we used Gaussian priors with means of 0. Participant-level parameters were drawn from group-level Gaussian distributions, the means, and precisions of which were again estimated from the data. We used Markov Chain Monte Carlo to estimate posterior distributions for all model parameters. Temporal discounting modeling was done in R ^[74]^ using the JAGS software package (version 4.3.0)^[75]^ and the Wiener module for JAGS ^[76]^. 2-step task modeling used Python in conjunction with the *pystan* toolbox and STAN (version 2.27) ^[77]^. We ran four chains for each model. For JAGS, chains consisted out of one million samples. Only the last 15000 were kept and the rest was discarded as a burn-in. For STAN, chains consisted of 7000 samples of which the first 3000 were discarded as a burn-in. Chain convergence was assessed using the 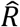 statistic^[78]^ where values ≤ 1.01 were considered acceptable. To determine the best fitting DDM in both tasks we calculated the Watanabe-Akaike Information Criterion (WAIC)^[79]^ using the loo-package (version 2.4.1) in R. Lower WAIC values represent better fits.

### Session and group effects on model parameters

To quantify potential session, group and interaction effects, we compared the group-level posterior distributions across groups and conditions. We then examined the 95% highest density intervals (HDI) and calculated directional Bayes Factors (dBF) quantifying the degree of evidence for a reduction vs. an increase in a parameter. Because priors for group-level parameters were symmetric, dBFs can simply calculated as the ratio of the posterior mass of the difference distribution above zero to the posterior mass below zero^[80]^. Here, dBFs above 3 are interpreted as moderate evidence in favor of a positive effect, while Bayes Factors above 12 are interpreted as strong evidence for a positive effect^[81]^. Specifically, a dBF of 3 would imply that a positive effect is three times more likely, given the data, than a negative effect. Bayes Factors below 0.33 are likewise interpreted as moderate evidence in favor of the alternative model with reverse directionality. A dBF above 100 is considered extreme evidence^[81]^. The cutoffs used here can be considered liberal, because they are usually used if the test is against a H_0_ implying an effect of exactly 0.

### Analysis of physiological data

Electrodermal activity (EDA) is a frequently employed index of sympathetic activity^[82]^. VR-related physiological arousal was first assessed using the tonic skin conductance level (SCL)^[62]^ via continuous deconvolution analysis (CDA), using default settings in the Ledalab toolbox^[83]^ for Matlab (The MathWorks). SCL was subsequently transformed into percentage change from the mean signal of the five-minute baseline phase and split into fifteen one-minute bins, five per experimental phase (pre-VR baseline [B], first exploration [F] and second exploration [S]). We additionally examined the number of spontaneous phasic skin conductance responses (SCR)^[83]^. The phasic component of the EDA signal was again divided into fifteen one-minute bins. For each bin we calculated the number of spontaneous SCRs. The resulting values were transformed into percentage change from the mean number of spontaneous SCRs during the five baseline bins.

Heart rate (HR) was analyzed in a similar fashion. The signal was converted into the percent change from baseline mean and divided into fifteen one-minute bins described above. Again, we compared bin B5 to F1 to assess the effect of immersion into VR and bins F5 and S1 to test for physiological cue-reactivity using Wilcoxon Signed-Ranks Tests^[84]^.

To test for general VR effects, the last baseline bin (B5) was compared to the first bin of the first exploration phase (F1) via non-parametric Wilcoxon Signed-Ranks Test^[84]^. To test for specific cue-reactivity effects, the last bin of the first exploration phase (F5) was compared to the first bin of the second exploration phase (S1), i.e. focusing on the time point were the specific gambling vs. neutral environment was entered, using non-parametric Wilcoxon Signed-Ranks Tests^[84]^. Effect sizes were calculated as the Z statistic divided by the square-root of N and reported as *r*^[85]^. Absolute r values above .5 are considered large effects, while values between .3 and .5 are considered medium. Values below .3 are considered small. The significance level was Bonferroni corrected for the number of statistical tests for each signal type.

### Subjective urge to gamble

Participants rated their subjective urge to gamble on a 10-point likert scale at five time points: at the beginning of the experiment (B1), at the end of the baseline phase (B5), at the end of the first exploration phase (F5), at the end of the second exploration phase (S5) and after completion of behavioral testing (end). Participants were prompted verbally and verbally reported their urge to gamble. Such verbal reports have been used successfully in previous cue-reactivity research ^[86]^ and for comparable VR designs^[41]^. Ratings were analyzed using a mixed model ANOVA.

## Results

### Subjective urge to gamble

Participants reported their subjective desire to gamble on a 10-point likert scale at five time points (see methods section). A mixed model ANOVA revealed a main effect of group (F(1) = 78.94, p <.001, η_p_^2^ = .434, i.e. an overall greater urge to gamble in the GD group) and a significant three-way interaction between group, session, and time point (F(3.059) = 2.81, p = .04, η_p_^2^ = .002). Simple effects analysis showed that this interaction was mostly due to an increased urge to gamble upon entering the gambling area in VR_gambling_ in the GD group but not the control group (t = -11.197, p < .001) (Figure 2). This increase was also significantly greater than the increase caused by the experimental area of the VR_neutral_ environment (t = -11.973, p < .001). VR_gambling_ exposure thus increased subjective craving specifically in the gambling group.

### Temporal discounting: model-agnostic analysis

A mixed model ANOVA on area-under-the curve values (AUC, a model-agnostic measure of temporal discounting, see methods section) revealed significantly steeper discounting (i.e., smaller AUC values) in gamblers vs. controls (F = 16.871, p <.001, η_p_^2^ = .216) (Figure 3 a and b). There was, however, no significant main effect of the VR session (F = .546, p = .463, η_p_^2^ < .001) and no significant group x session interaction (F = 1.686, p = .199, η_p_^2^ = .002).

**Figure 3.**
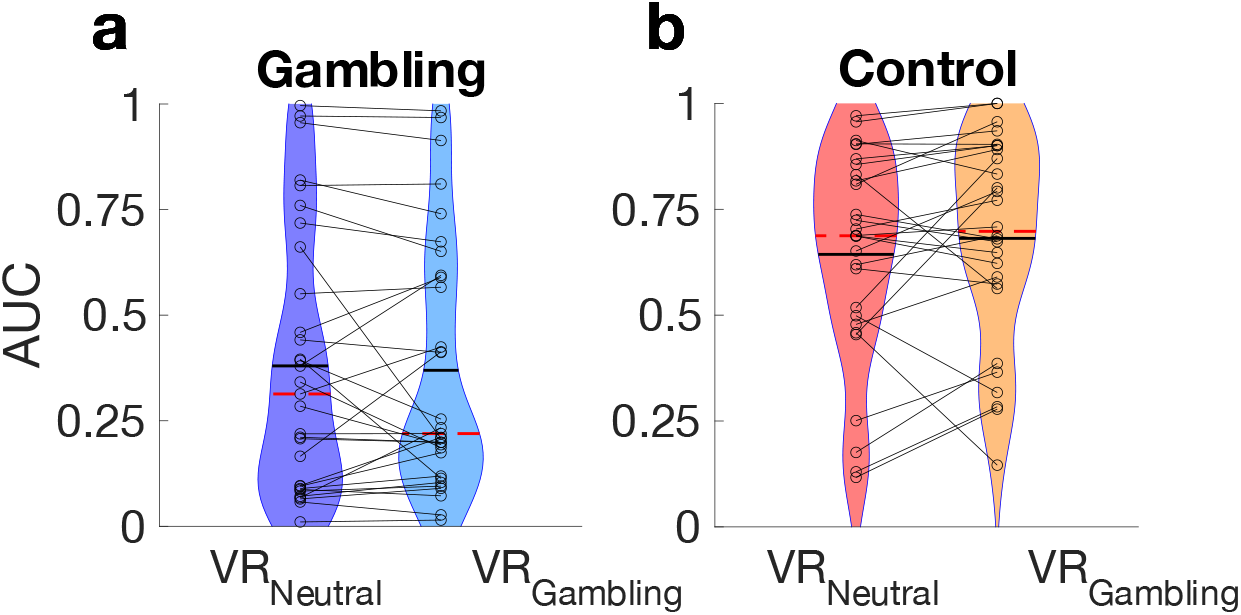
Violin plot of AUC values for the GD group (a) and non-gambling controls (b). Lines connect single participant data points. Dashed red lines represent the median and solid black lines the mean.

### Temporal discounting softmax choice rule

Group level posteriors revealed extreme evidence for an increased discount rate log(k) in the gambling group (dBF>100, 95% HDI: min = .52, max = 2.501) (Figure 4 a and Table 2). However, there was only moderate evidence a further increase in discounting in the VR_gambling_ environment gamblers (s_k_ dBF = 3.338; 95% HDI: min = -.156, max = .325) (Figure 4 c and Table 3). In contrast, controls showed strong evidence for reduced discounting in the VR_gambling_ session (dBF = 0.072, 95% HDI: min = - .733, max = .081). Decision noise (*β*) was overall lower in controls vs. gamblers (dBF <0.01, 95% HDI: min = -.492, max = -.08) (Figure 4 b and Table 2). There was no conclusive evidence for a VR effect on *β* in either group (Controls: dBF = 1.410, 95% HDI: min = -.055, max = .065; Gambling: dBF = 2.476, 95% HDI: min = -.087, max = .17) (Figure 4 d and Table 3).

**Figure 4.**
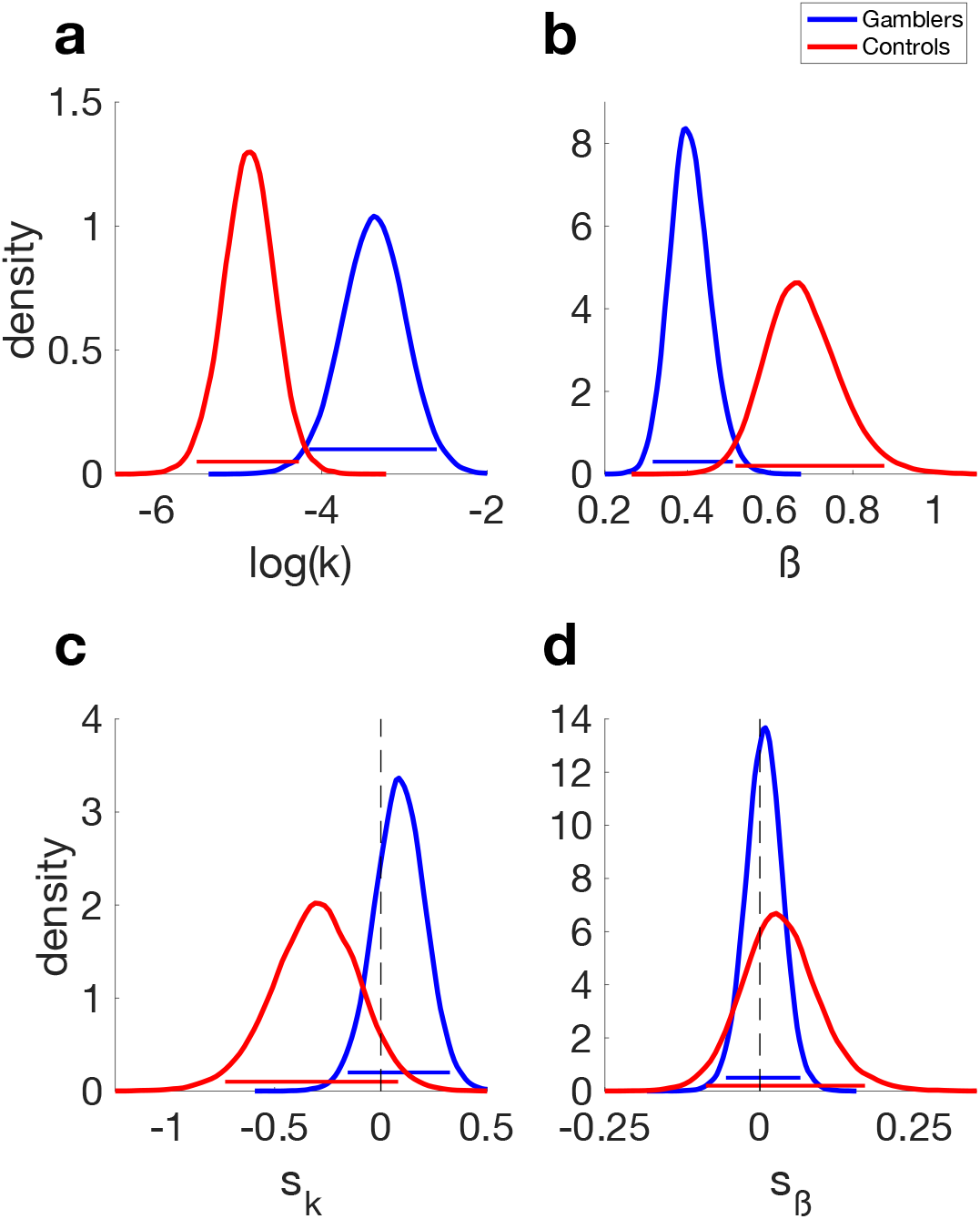
Hyperbolic temporal discounting model with a softmax choice rule. Group level posterior distributions for the gambling (blue) and control group (red). a) discount-rate log(k). b) softmax *β*. c) shift parameter s_k_ describing the shift in log(k) from the VR_neutral_ to the VR_gambling_ session. d) shift parameter *s*_*β*_ describing the shift in softmax *β* from the VR_neutral_ to the VR_gambling_ session. Horizontal lines denote 95% highest posterior density intervals.

**Table 2.** Posterior means and directional Bayes Factors (dBF) for the parameters of the hyperbolic temporal discounting model with softmax (left) and DDM_S_ choice rule (right). dBF values around 1 indicate that values are evenly distributed around 0. dBFs are calculated as BF = i/(1-i), with i being the probability mass of the posterior distributions above zero. dBFs for group difference are based on the difference distributions between groups. Values are reported as Gambling > Control.

| Model parameter | Softmax Model |  |  | DDMs |  |  |  |  |
| --- | --- | --- | --- | --- | --- | --- | --- | --- |
|  | GD: Mean | CON: Mean | dBF group | GD: Mean | CON: Mean | dBF group | GD: dBF vs. 0 | CON: dBF vs. 0 |
| $\log(k)$ | -3.371 | -4.884 | 448.116 | -3.428 | -4.791 | 353.78 | - | - |
| $\beta$ | .406 | .679 | .003 | - | - | - | - | - |
| $\nu_{coeff}$ | - | - | - | .21 | .267 | .106 | >100 | >100 |
| $\nu_{max}$ | - | - | - | 1.772 | 1.86 | .387 | >100 | >100 |
| $\alpha$ | - | - | - | 2.85 | 3.013 | .224 | - | - |
| $z$ | - | - | - | .499 | .544 | .01 | - | - |
| $\tau$ | - | - | - | .911 | 1.09 | .023 | - | - |

**Table 3.** Mean and Directional Bayes Factors (dBF) values for the shift parameters of the hyperbolic temporal discounting model with a softmax and a DDMs choice rule. dBFs values around 1 indicate that values are evenly distributed around 0. dBFs are calculated as BF = i/(1-i), with i being the probability mass of the posterior distributions above zero. dBFs for group difference are based on the difference distributions between groups. Values are reported as Gambling > Control. dBFs for group comparisons are based on the difference distributions of the posteriors of both groups.

| Model parameter (shift) | Softmax Model |  |  |  |  | DDMs |  |  |  |  |
| --- | --- | --- | --- | --- | --- | --- | --- | --- | --- | --- |
|  | GD: Mean | CON: Mean | dBF group | GD: dBF vs. 0 | Con: dBF vs. 0 | GD: Mean | CON: Mean | dBF group | GD: dBF vs. 0 | Con: dBF vs. 0 |
| $s_k$ | .088 | -.309 | 19.457 | 3.338 | .072 | -.01 | -.442 | 21.888 | 1.119 | .031 |
| $s_\beta$ | .006 | 2.476 | .564 | 1.41 | 2.476 | - | - | - | - | - |
| $s_{vcoeff}$ | - | - | - | - | - | .03 | .096 | .137 | 6.643 | 48.758 |
| $s_{vmax}$ | - | - | - | - | - | .047 | .104 | .516 | 3.499 | 4.675 |
| $s_\alpha$ | - | - | - | - | - | -.196 | -.029 | .263 | .089 | .707 |
| $s_z$ | - | - | - | - | - | .001 | .007 | .615 | 1.195 | 2.166 |
| $s_\tau$ | - | - | - | - | - | .02 | .065 | .369 | 1.577 | 6.964 |

### Temporal discounting drift diffusion model choice rule

Model comparison revealed that the DDM_S_ (DDM including a sigmoidal drift rate modulation) accounted for the data best (Table 4), replicating previous findings^[39,46,48,51]^. Further analyses focused on the DDM_S_. The group-level posterior distributions for log(k) again indicated extreme evidence for stronger discounting in the gambling group (dBF > 100, 95% HDI: min = .43, max = 2.293) (Figure 5 a and Table 2). There was only anecdotal evidence for an increase in temporal discounting in the VR_gambling_ session (modelled by the s_k_ parameter) in the gambling group (dBF = 1.119, 95% HDI: min = -.234, max = .249) (Figure 6 a and Table 3). In the control group however, there was strong evidence for decreased discounting in the VR_gambling_ session (dBF = 0.031, 95% HDI: min = -.915, max = .004) (Figure 6 a and Table 3). For the v_coeff_ parameter, mapping trial wise value differences to the drift rate, we found moderate evidence in favor of a higher influences of trial wise value differences in the control group compared to the gambling group (dBF = .106, 95% HDI: min = -.144, max = .029) (Figure 5 d and Table 2), mirroring the effects observed for the decision noise parameter *β* in the softmax model. The gambling and the control groups displayed moderate and very strong evidence in favor of an increase in v_coeff_ in the VR_gambling_ session (*s*_*νcoeff*_ Gambling: dBF = 6.643, 95% HDI: min = -.019, max = .087; Control: dBF = 48.758, 95% HDI: min = .007, max = .2) (Figure 6 d and Table 3). For *ν*_max_, the upper boundary for the trial-wise value difference’s influence on the drift rate, the dBF indicated only anecdotal evidence for a difference between groups (dBF = 0.387, 95% HDI: min = -.37, max = .193) (Figure 5 b and Table 2). Furthermore, both groups showed moderate evidence for an increase in *ν*_max_ in the VRgambling session (s*ν*_max_ Gambling: dBF = 3.499, 95% HDI: min = -.073, max = .167; Control: dBF = 4.675, 95% HDI: min = -.113, max = .325) (Figure 6 b and Table 3). The group level posterior distributions for the *α* parameter, governing the trade-off between speed and accuracy, showed only moderate evidence in favor of an increased accuracy focus in controls (dBF = 0.224, 95% HDI: min = -.524, max =.194) (Figure 5 e and Table 2). s_*α*_ parameter group level posterior distributions revealed strong evidence for a decrease in boundary separation in VR_gambling_ in gamblers (dBF = 0.089, 95% HDI: min = - .478, max = .089) (Figure 6 e and Table 3). For controls, there was no conclusive evidence for directional effects (dBF = 0.707, 95% HDI: min = -.313, max = .259) (Figure 6 e and Table 3). The bias parameter *z* revealed extreme evidence for a more pronounced bias towards SS vs. LL choices in gamblers vs. controls (dBF = 0.009, 95% HDI: min = -.083, max = -.008) (Figure 5 c and Table 2). Session effects on *z* were of inconclusive directionality (s_*z*_ Gambling: dBF = 1.195, 95% HDI: min = -.024, max = .026; Control: dBF = 2.166, 95% HDI: min = -.02, max = .034) (Figure 6 c and Table 3). For the non-decision time *τ* there was strong evidence for a reduced value in gamblers (dBF = 0.023, 95% HDI: min = -.381, max = -.005), reflecting faster motor and/or perceptual RT components (Figure 5 f and Table 2). There was moderate evidence for an increased non-decision time in the control group in the VRgambling session (s*τ* dBF = 6.924, 95% HDI: min = -.094, max = .135). This was not the case for the gambling group (s*τ* dBF = 1.576, 95% HDI: min = -.045, max = .175) (Figure 6 f and Table 3).

**Table 4.** Summary of the WAICs of all drift diffusion models (DDM) in all sessions. Ranks are based on the lowest WAIC in all sessions.

| Model | Gambling |  | Controls |  | Rank |
| --- | --- | --- | --- | --- | --- |
|  | VR<sub>neutral</sub> | VR<sub>gambling</sub> | VR<sub>neutral</sub> | VR<sub>gambling</sub> |  |
| DDM<sub>0</sub> | 8197.4 | 7563.7 | 7558.3 | 6925.3 | 3 |
| DDM<sub>L</sub> | 6596.5 | 6021.7 | 6296.5 | 5748 | 2 |
| DDM<sub>S</sub> | 6243.5 | 5611 | 5619 | 5142.4 | 1 |

**Figure 5.**
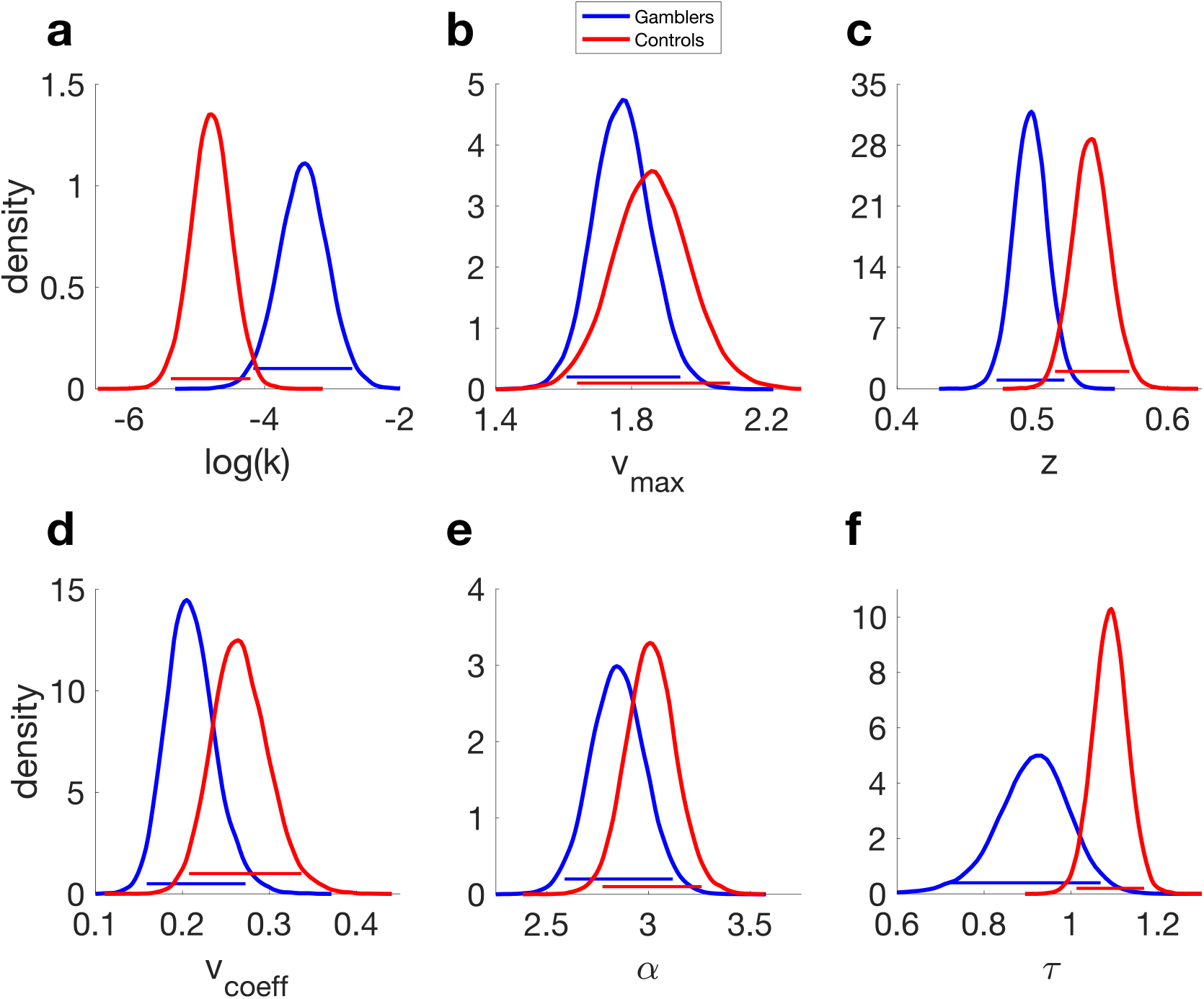
Hyperbolic temporal discounting model with drift diffusion model choice rule: parameter posterior distributions from the VR_neutral_ condition. a) discount-rate log(k). b) maximum drift-rate v_max_. c) starting-point z. d) drift-rate coefficient v_coeff_. e) boundary separation α. f) non-decision time *τ*. Horizontal lines denote 95% highest posterior density intervals.

**Figure 6.**
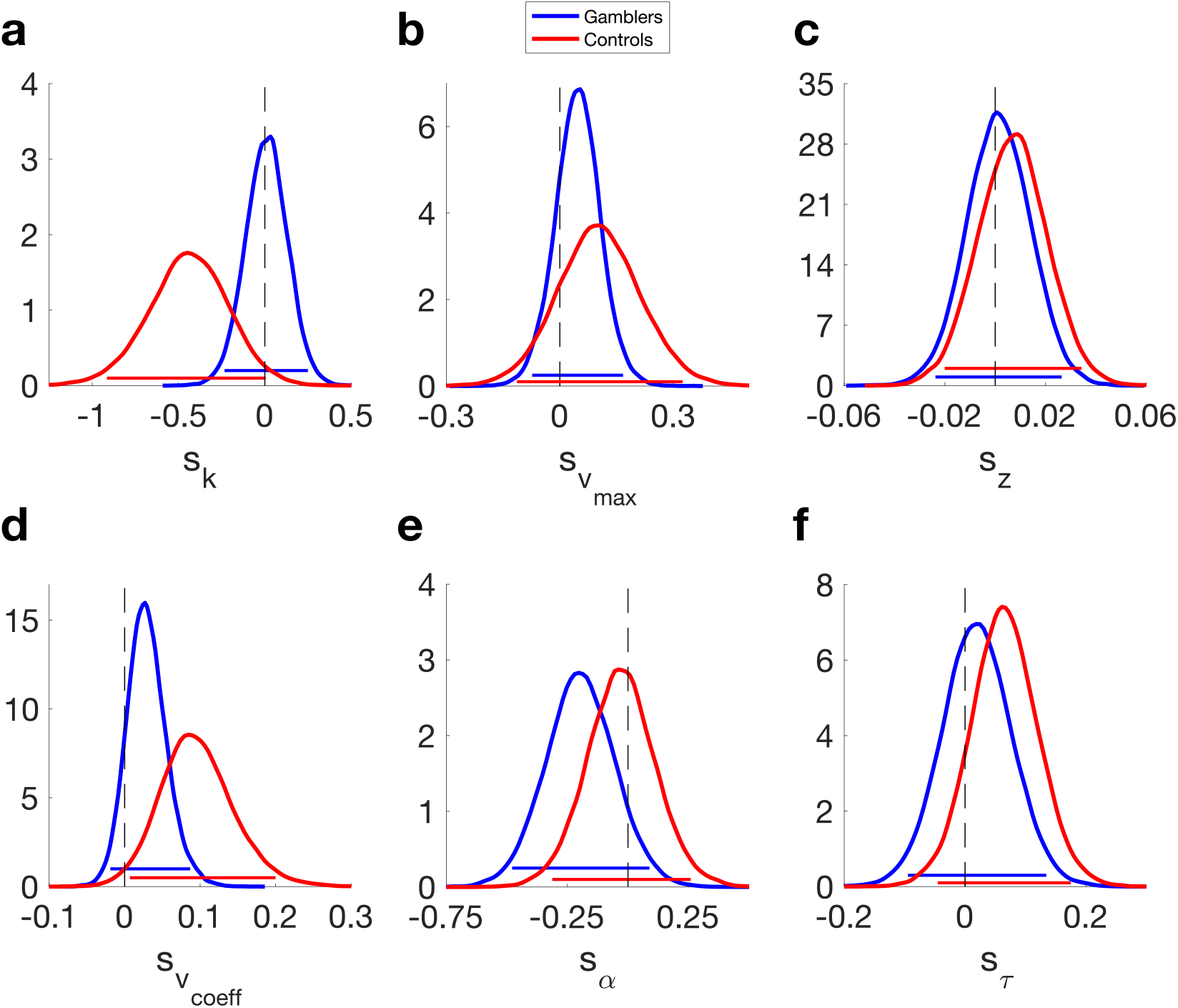
Hyperbolic temporal discounting model with drift diffusion model choice rule: s-parameters modeling the change in each parameter from the VR_neutral_ to the VR_gambling_ session. a) discount-rate log(k). b) maximum drift-rate v_max_. c) starting-point z. d) drift-rate coefficient v_coeff_. e) boundary separation α. f) non-decision time *τ*. Horizontal lines denote 95% highest posterior density intervals.

### 2-step task: model-agnostic analyses

Numerically, non-gambling controls scored more points overall (Mean [SD]: Gambling group, VR_gambling_: 118.835 [9.931], VR_neutral_: 116.751 [10.768], Control group, VR_gambling_: 120.096 [11.066], VR_neutral_: 121.535 [10.524]), although a mixed model ANOVA revealed that the main effect of group did not reach significance (F(1) = 3.788, p = .056, η_p_^2^ = .021). Likewise, there was no significant effect of session (F(1) = .021, p = .887, η_p_^2^ = <.001), nor a group x session interaction (F(1) = .615, p = .436, η_p_^2^ = .007).

Second stage RTs were then analyzed as a function of transition type (common vs. rare), session (VR_gambling_ vs. VR_neutral_) and group (gambling vs. non-gambling) using a hierarchical linear mixed model (see Figure 7 a to d). Here, increases in S2 RTs following rare transitions are taken as a measure of MB control^[70]^. This revealed significant effects of transition (p < .001, see Table 5), session (p = .002, see Table 5) and group (p < .001, see Table 5) as well as a significant interaction between the transition and group (p < .001, see Table 5), reflecting a greater increase in S2 RTs following rare transitions in controls. A further model-agnostic analysis of the 2-step task typically entails an analysis of S1 stay probabilities (i.e. the probability of repeating the S1 choice made on the previous trial) as a function of previous reward and transition^[14]^. Since in the 2-step task version employed here, S2 rewards were not probabilistic but continuous we ran an analogous analysis using a moving average of recent rewards to categorize trials according to previous rewards. Results are presented in the supplementary materials (Supplementary Table 3). Resonating with the results from the RT analysis, this analysis revealed MF and MB influences on S1 decisions in both groups (the factor Reward and the interaction of the factors Reward*Transition both were significant at p < .001, Supplementary Table 3). Additionally, we observed a greater MB effect in the non-gambling group (significant three-way interaction of Reward*Transition*Group at p < .001, Supplementary Table 3).

**Figure 7.**
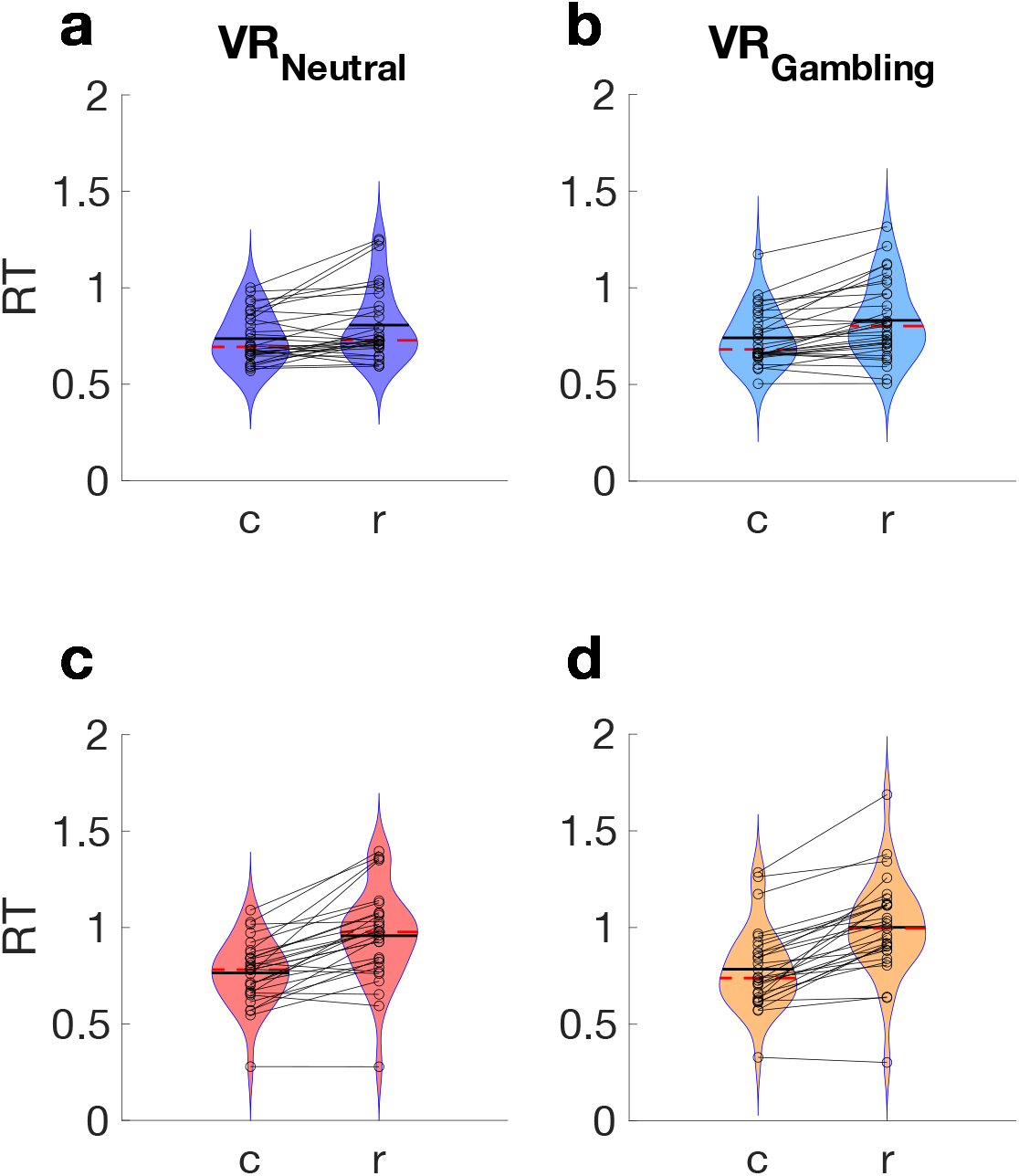
Violin plots of median S2 response times (RTs) from the 2-step task per group (top row: gambling group, bottom row: non-gambling group) and condition. Lines connect individual participant data points. a) gambling group, VR_neutral_, b) gambling group, VR_gambling_. c) control group, VR_neutral_. d:) control group, VR_gambling_. “c” denotes common transitions, “r” denotes rare transitions. Dashed red lines represent the median and solid black lines the mean.

**Table 5.** Results of the Hierarchical General Linear Model analysis of second stage RTs from the 2-step task, with Transition (common vs. rare), Session (gambling vs. neutral) and Group (Gambling vs. Non-gambling) as fixed effects and subject as random effect.

| Fixed effects |  |  |  |  |
| --- | --- | --- | --- | --- |
|  | Estimate | Std. Error | *t* | *p*-value |
| **Intercept** | -.067 | .026 | -2.61 | **.013** |
| **Transition** | -.183 | .01 | -17.761 | **<.001** |
| **Session** | .037 | .012 | 3.043 | **.002** |
| **Group** | -.157 | .012 | -13.038 | **<.001** |
| **Trans\*Session** | -.022 | .015 | -1.494 | .135 |
| **Trans\*Group** | 1.121 | .014 | 8.46 | **<.001** |
| **Session\*Group** | -.015 | .017 | -.888 | .375 |
| **Trans\*Session\*Group** | -.004 | .02 | -.179 | .858 |

### 2-step task: softmax choice rule

As pre-registered, we initially applied the hybrid model proposed by Daw and colleagues^[14]^ and extended by Otto et al.^[71]^. The model includes independent learning rates for S1 and S2, MF and MB *β* weights for S1, and a single softmax slope parameter *β* for S2 (see methods). In both groups, S1 choices were modulated by MF Q-values (*β*_*MF*_ Gambling: dBF >100, 95% HDI: min = 1.504, max = 3.247, Controls: dBF >100, 95% HDI: min = 1.325, max = 2.682) (Figure 8 c) and MB Q-values (*β*_*MB*_ Gambling: dBF = 64.204, 95% HDI: min = .245, max = 5.379, Controls: dBF >100, 95% HDI: min = 4.505, max = 9.597) (Figure 8 b). *β*_*MB*_ was greater in controls (dBF = .011, 95% HDI: min = -7.791, max = -.64), but this was not the case for *β*_*MF*_ (BFs: 2.988, 95% HDI: min = -.757, max = 1.473). There was only moderate evidence for a gambling-session related increase in *β*_*MB*_ in both groups (Gambling: dBF: 4.692, 95% HDI: min = -.922, max = 2.811, Control: dBF: 8.442, 95% HDI: min = -.657, max = 3.455) (Figure 8 g). Other parameters modeling VR_gambling_ effects only revealed if anything anecdotal evidence (see Figure 8 a to j and Tables 6 and 7). For a full list of all directional Bayes Factors please refer to Tables 6 and 7.

**Figure 8.**
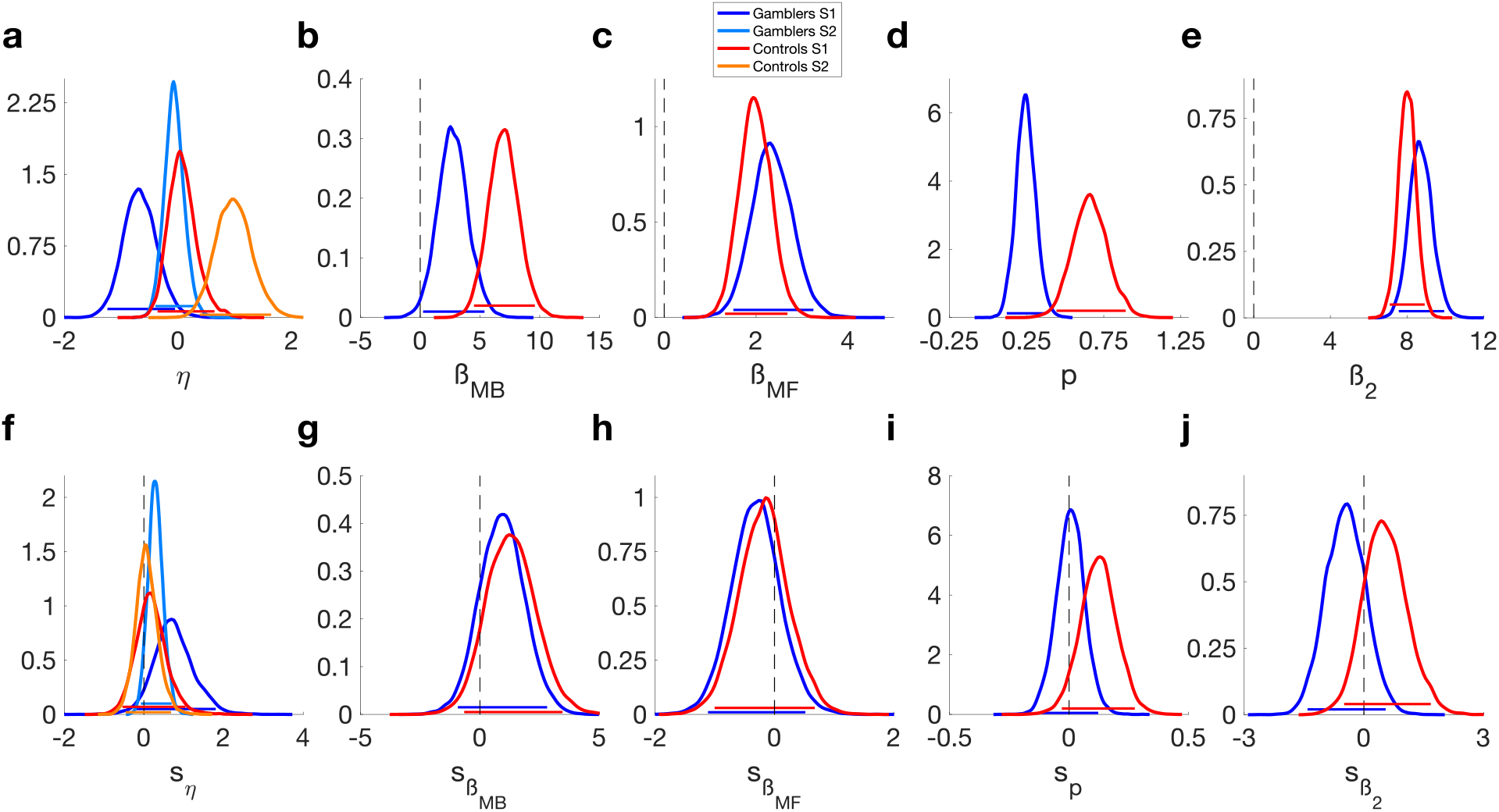
Hybrid model with softmax choice rule posterior distributions. Top row: Parameters values for the VR_neutral_ session, Bottom row: shifts in parameter values in VR_gambling_ session. a) First and second stage learning rates. b) MB weights *β*_*MB*_. c) MF weights *β*_*MF*_. d) perseverance parameter p. e) Second stage softmax *β*_2_. f) shift in first and second stage learning rates. g) shift in *β*_*MB*_. h) shift in *β*_*MF*_. i) shift in p. j) shift in *β*_2_. Horizontal lines denote 95% highest posterior density intervals.

**Table 6.** Mean and Directional Bayes Factors (dBF) values for the parameters of the Hybrid Rl model with a softmax and a DDMs choice rule. dBFs values around 1 indicate that values are evenly distributed around 0. dBFs are calculated as BF = i/(1-i), with i being the probability mass of the posterior distributions above zero. dBFs for group difference are based on the difference distributions between groups. Values are reported as Gambling > Control. dBFs for group comparisons are based on the difference distributions of the posteriors of both groups.

| Model Parameter | Softmax Model |  |  |  |  | DDMs |  |  |  |  |
| --- | --- | --- | --- | --- | --- | --- | --- | --- | --- | --- |
|  | GD: Mean | Con: Mean | dBF group | GD: dBF vs. 0 | Con: dBF vs. 0 | GD: Mean | Con: Mean | dBF group | GD: dBF vs. 0 | Con: dBF vs. 0 |
| $\eta_1$ | .749 | .188 | .034 | - | - | -.355 | -.282 | .818 | - | - |
| $\eta_2$ | .293 | .075 | .002 | - | - | .012 | .62 | .057 | - | - |
| $\beta_{MB}$ | 2.744 | 6.981 | .011 | 64.204 | >100 | - | - | - | - | - |
| $\beta_{MF}$ | 2.348 | 1.985 | 2.988 | >100 | >100 | - | - | - | - | - |
| $\beta_2$ | 8.718 | 8.014 | 4.503 | >100 | >100 | - | - | - | - | - |
| $p$ | .245 | .662 | <.001 | - | - | .311 | .708 | .011 | - | - |
| $v_{coeff\ MB}$ | - | - | - | - | - | 3.17 | 8.895 | .009 | 69.665 | >100 |
| $v_{coeff\ MF}$ | - | - | - | - | - | 2.959 | 2.271 | 3.802 | >100 | >100 |
| $v_{coeff2}$ | - | - | - | - | - | 6.87 | 6.482 | 1.869 | >100 | >100 |
| $v_{max1}$ | - | - | - | - | - | .567 | 1.26 | .185 | 30.118 | >100 |
| $v_{max2}$ | - | - | - | - | - | 2.491 | 2.707 | .252 | >100 | >100 |
| $\alpha_1$ | - | - | - | - | - | 1.358 | 1.427 | .22 | - | - |
| $\alpha_2$ | - | - | - | - | - | 1.527 | 1.587 | .162 | - | - |
| $\tau_1$ | - | - | - | - | - | 1.738 | 1.896 | .002 | - | - |
| $\tau_2$ | - | - | - | - | - | .391 | .421 | .196 | - | - |

**Table 7.**
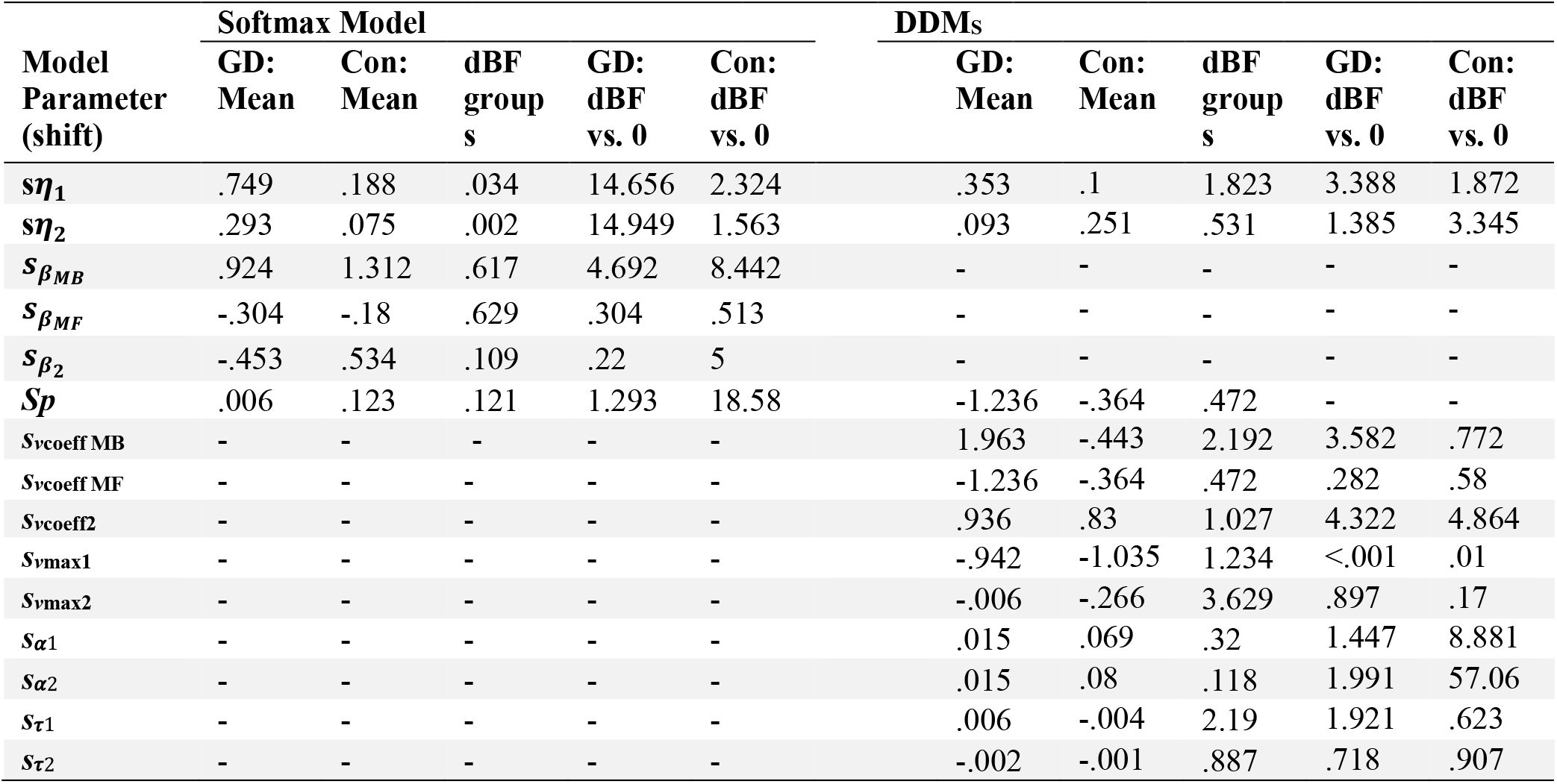
Posterior means and Directional Bayes Factors (dBF) for parameters modeling condition effects for 2-step task models (left: hybrid RL model softmax choice rule, right: hybrid RL model with DDM_S_ choice rule). dBF values around 1 indicate that values are evenly distributed around 0. dBFs are calculated as BF = i/(1-i), with i being the probability mass of the posterior distributions above zero. dBFs for group difference are based on the difference distributions between groups. Values are reported as Gambling > Control.

### 2-step task: drift diffusion model choice rule

Next, we replaced the softmax choice rule with the DDM. In line with the results from the temporal discounting task, a non-linear drift-rate scaling accounted for the data best, and this was the case in both groups (Table 8). In both groups, S1 choices were affected by both MF Q-value differences (*ν*_coeffMF_ > 0, Gambling: dBF >100, 95% HDI: min = 1.828, max = 4.3; Control: dBF >100, 95% HDI: min = 1.34, max = 3.408) (Figure 9 f and Table 6) and MB Q-value differences (*ν*_coeffMB_ > 0, Gambling: dBF = 69.665, 95% HDI: min = .414, max = 6.363; Control: dBF >100, 95% HDI: min = 5.638, max = 12.665) (Figure 9 e and Table 6). As in the softmax model, we observed extreme evidence for a greater MB effect (*ν*_coeffMB_) in the non-gambling control group (dBF:.009, 95% HDI: min = -10.483, max = -1.159) (Figure 9 e and Table 6). For MF Q-values we observed moderate evidence for a higher *ν*_coeffMF_ ion the gambling group compared to the non-gambling control group (BFs: 3.802, 95% HDI: min = -.915, max = 2.346) (Figure 9 f and Table 6). In both groups, we there was only anecdotal or inconclusive evidence for parameter changes in the VR_gambling_ session (Figure 10 a to h and Table 7).

**Table 8.**
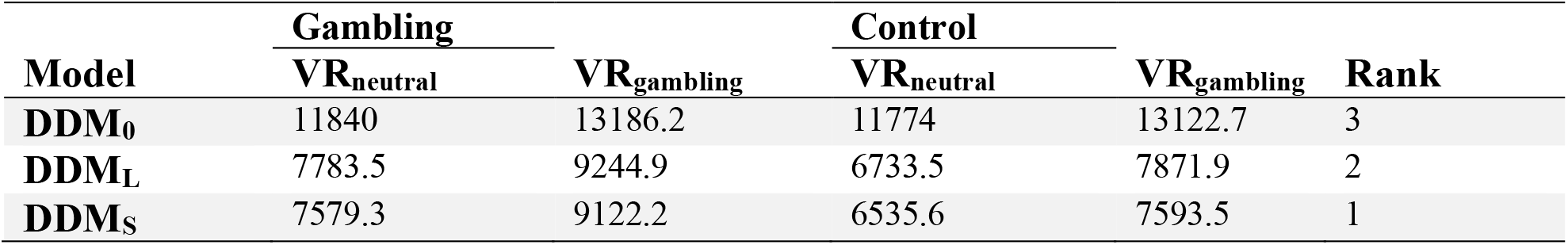
Summary of the WAICs of all DDM models in all sessions. Ranks are based on the lowest WAIC in all sessions.

**Figure 9.**
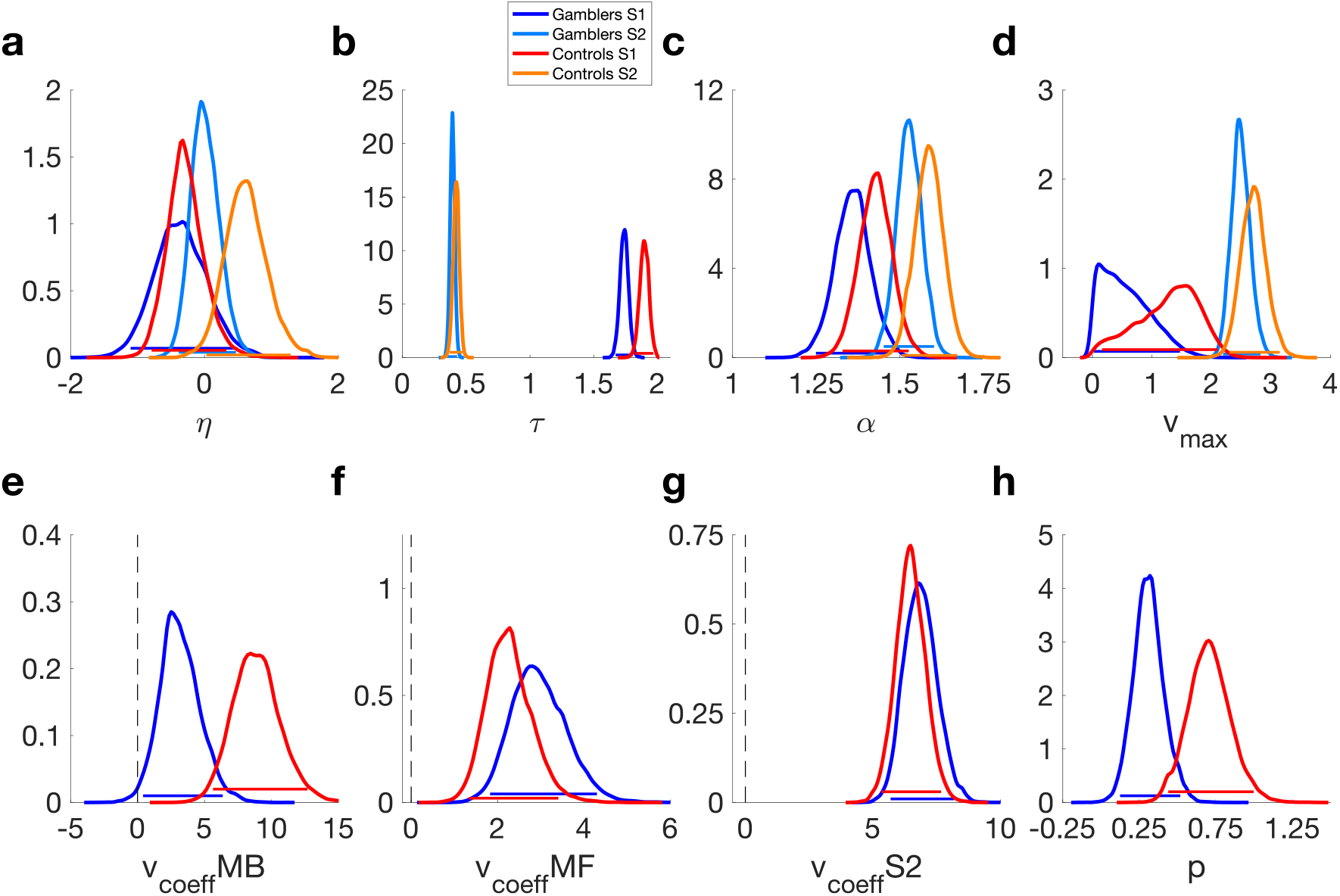
Hybrid RL model with a non-linear drift diffusion model choice rule. a) learning rates for S1 and S2 *η*. b) non-decision time *τ*. c) boundary separation α. d) maximum drift-rate v_max_. e) MB drift-rate coefficient v_coeffMB_. f) MF drift-rate coefficient v_coeffMF_. g) drift-rate coefficient v_coeff_ for S2. h) perseverance parameter p. Horizontal lines denote 95% highest posterior density intervals.

**Figure 10.**
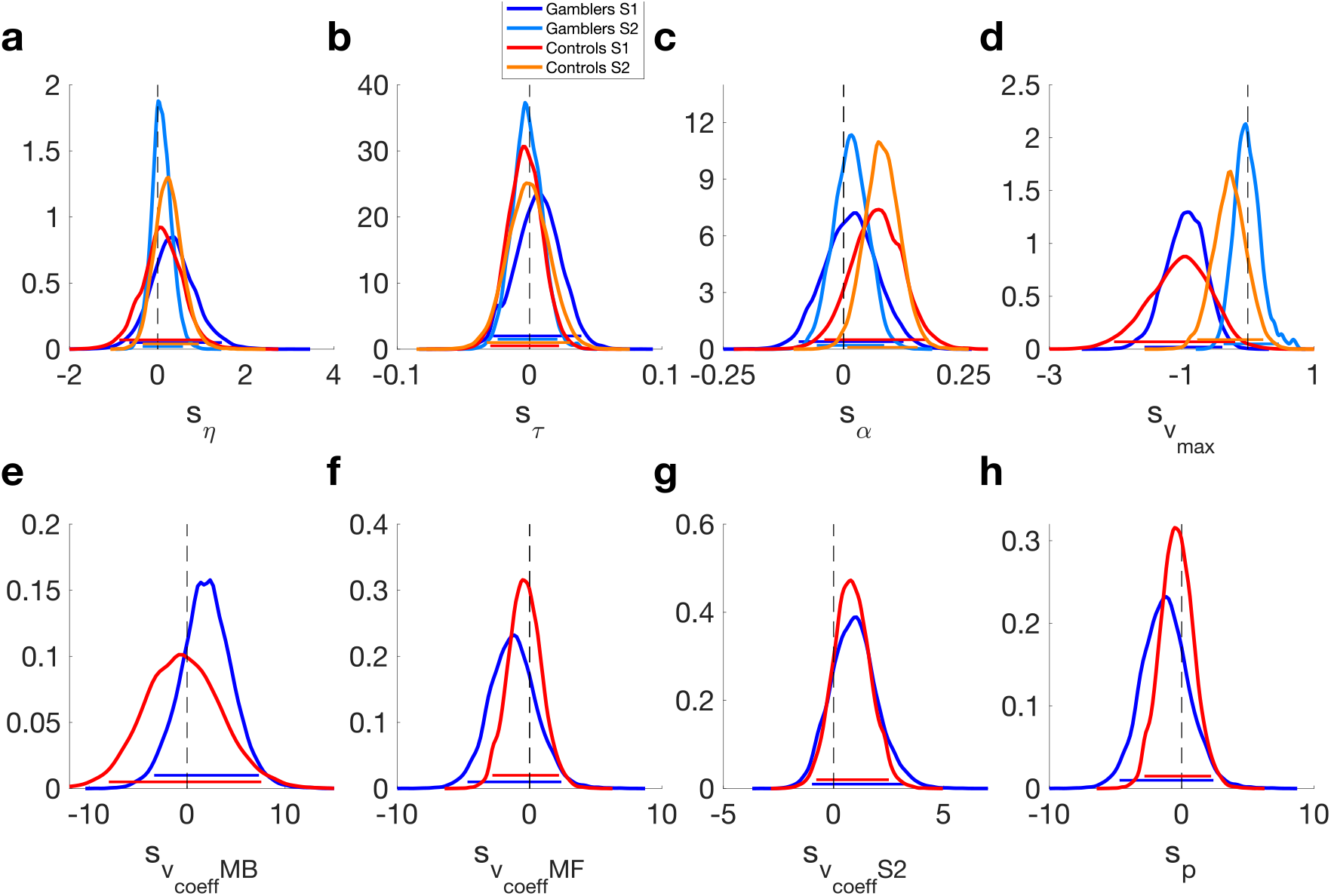
Posterior distributions of parameters modeling condition effects for the hybrid RL model with drift diffusion model choice rule (DDM_S_). a) shift in the learning rate for S1 and S2 *s*_*η*_. b) shift in non-decision time s_*τ*_. c) shift in boundary separation s_α_. d) shift in maximum drift-rate s_vmax21_. e) shift in MB drift-rate coefficient s_vcoeffMB_. f) shift in MF drift-rate coefficient s_vcoeffMF_. g) shift in drift-rate coefficient s_vcoeff_ for S2. h) shift in perseverance parameter s_p_. Horizontal lines denote 95% highest posterior density intervals.

### Electrodermal activity (EDA)

As preregistered, psychophysiological cue-reactivity was analyzed by converting the number of spontaneous skin conductance responses (nSCR) and the skin conductance level (SCL) into percentage change from baseline. Per phase, values were binned into fifteen one-minute intervals (five each for the baseline phase [B], first exploration phase [F] and second exploration phase [S]). All comparisons were tested for significance with the Wilcoxon Signed Rank Test. The significance level was Bonferroni corrected. Entering VR (i.e., B5 vs. F1) led to a significant increase in SCL in both groups and there was no significant difference between conditions (Figures 11 and 12 c and d and Supplementary Tables 4 and 5). The effect size was large throughout (r > .5). There was no corresponding significant change in nSCRs in either group (Figures 11 and 12 a and b Supplementary Tables 6 and 7). Psychophysiological cue-reactivity was examined by comparing the difference between F5 and S1 (i.e. the effect of entering the experimental areas: virtual café vs. virtual casino) between VR_neutral_ and VR_gambling_ per group. There were no significant effects on either nSCR or SCL and therefore no evidence for psychophysiological cue-reactivity in nSCRs (Figure 11 and 12 a and b and Supplementary Tables 6 and 7) and SCL (Figures 11 and 12 c and d and Supplementary Tables 4 and 5).

**Figure 11.**
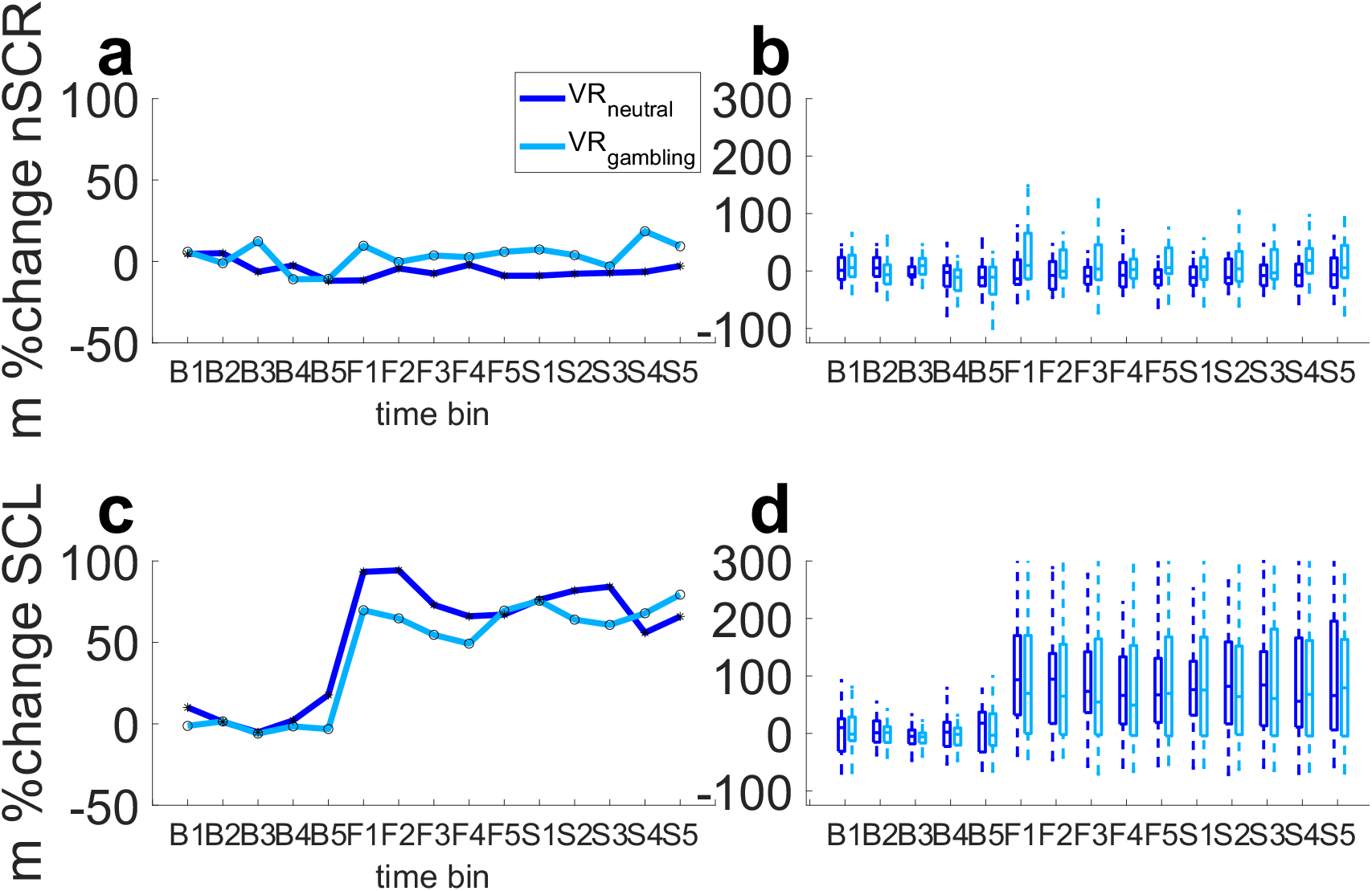
Results of the EDA measurements in the gambling group divided into 15 time points over the course of the baseline phase, measured before participants entered the VR-environments, and the first and second exploration phases. Each of the three phases is divided into five one-minute bins (B1-5: pre-VR baseline, F1-5: first exploration phase in VR, S1-5: second exploration phase VR). a: Median percent change from baseline mean for no. of spontaneous SCRs over gambling participants. b: Boxplot of percentage change from baseline mean for no. spontaneous SCRs over gambling participants. c: Median percent change from baseline mean of SCL over gambling participants. d: Boxplots of percentage change from base line mean of SCL over gambling participants.

**Figure 12.**
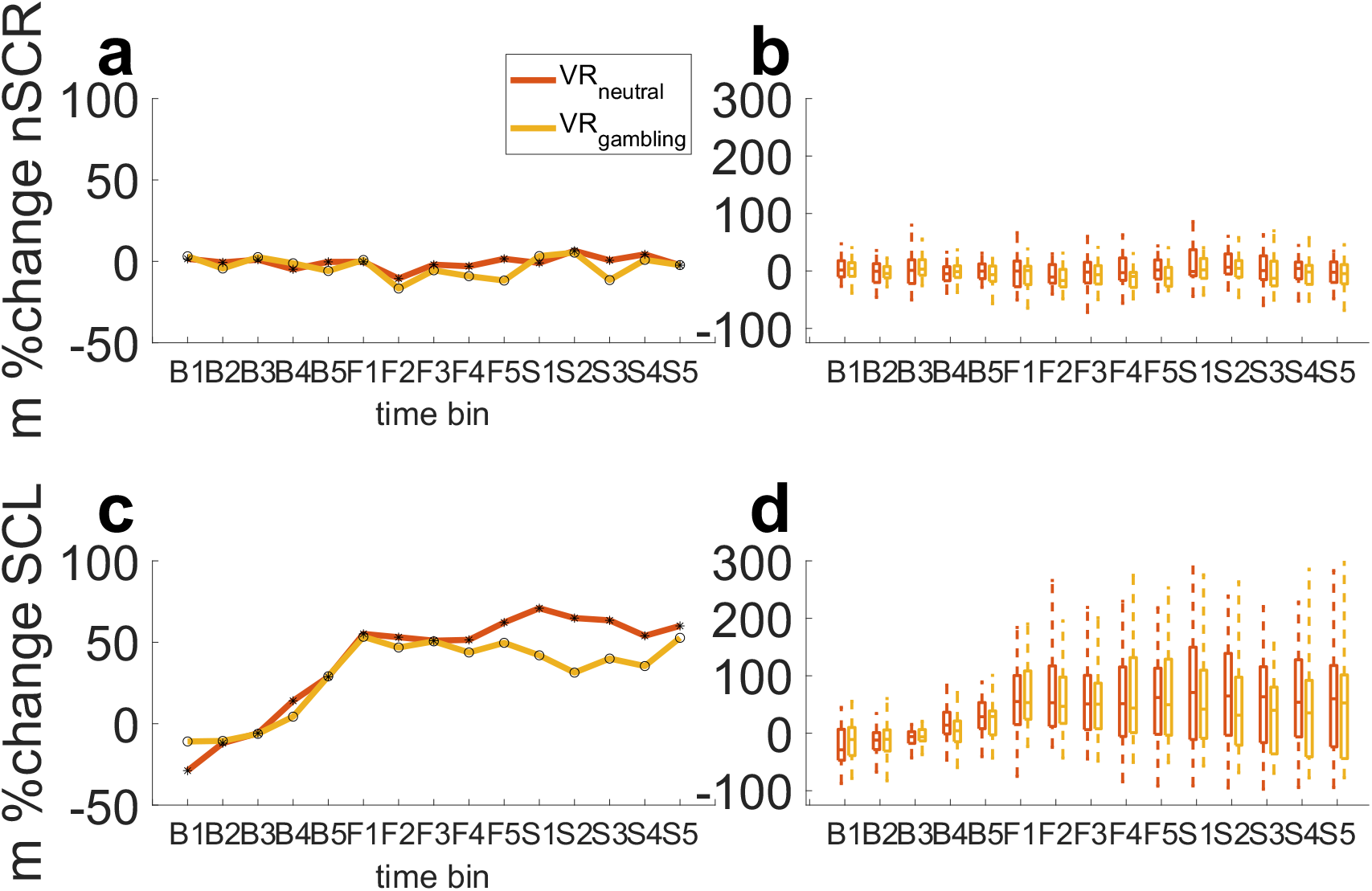
Results of the EDA measurements in the control group divided into 15 time points over the course of the baseline phase, measured before participants entered the VR-environments, and the first and second exploration phases. Each of the three phases is divided into five one-minute bins (B1-5: pre-VR baseline, F1-5: first exploration phase in VR, S1-5: second exploration phase VR). a: Median percent change from baseline mean for no. of spontaneous SCRs over control participants. b: Boxplot of percentage change from baseline mean for no. spontaneous SCRs over control participants. c: Median percent change from baseline mean of SCL over control participants. d: Boxplots of percentage change from base line mean of SCL over control participants.

### Heart Rate (HR)

Analysis of HR proceeded along similar lines as the analysis of the EDA data described above. HR was first converted into percent signal change from baseline, and then divided into fifteen one-minute bins (five each for the baseline phase [B], first exploration phase [F] and second exploration phase [S]). We observed no overall significant increase in HR in response to VR immersion (B5 vs. F1) in either group and environment, with the VR_gambling_ environment in the gambling group forming the single exception (Supplementary Tables 8 and 9). However, this effect was not significantly greater than in VR_neutral_. Likewise, entering the experimental areas of the VR environments did not significantly increase HR (Supplementary Tables 8 and 9) in either group or environment. Accordingly, there was no evidence for significant cue-reactivity effects on HR.

## Discussion

Here we investigated the subjective, behavioral, and physiological effects of virtual reality (VR) gambling environment exposure in regular gamblers (GD group) and matched non-gambling controls. The joint assessment of these three levels of cue-reactivity in VR enabled us to thoroughly delineate several possible effects, thereby informing potential future application of VR in addiction research. Participants explored two rich and navigable virtual environments (a café environment and a casino/sports betting environment: VR_neutral_ vs. VR_gambling_) and within both environments performed two behavioral tasks with high relevance for gambling disorder and addiction, temporal discounting, and a 2-step sequential RL task. In both groups, exposure to VR substantially increased sympathetic arousal as reflected in the tonic skin conductance level (SCL). However, despite the fact that the VR_gambling_ environment selectively increased subjective craving in the GD group, no physiological measure showed a pattern consistent with physiological cue-reactivity in the gambling group. Analysis of the behavioral data revealed that previously observed group differences between gamblers and controls were replicated in VR. First, gamblers discounted delayed rewards more steeply than controls^[1,5,9,10]^, but this effect was not differentially modulated by the VR_gambling_ environment. Second, gamblers relied less on a model-based (MB) decision strategy during the 2-step sequential RL task^[17]^, but this effect was again not differentially modulated by the VR_gambling_ environment.

### Self-reported cue-reactivity

Participants verbally reported subjective urge-to-gamble (craving) at four time points during exposure to the virtual environments. Craving was overall higher in the gambling group, and exposure to the VR_gambling_ environment selectively increased the subjective craving in gamblers but not controls. This is in line with earlier results^[40]^ showing that virtual gambling environments induce subjective craving in frequent gamblers on a level comparable to gambling on real video slot machines. Thus, our VR environment exhibited ecological validity with respect to self-reported urge-to-gamble in gambling participants.

### Physiological cue-reactivity

How addiction manifests on a computational, physiological and neural level has important implications for treatment and relapse prevention. Alterations in neural reward processing in ventral striatum and ventromedial prefrontal cortex have been frequently observed in gambling disorder, albeit with considerable heterogeneity in the directionality of these effects^[87]^. Gambling-related visual cues might interfere with striatal valuation signals in GD, and might in turn increase temporal discounting^[6]^. Here, analysis of physiological cue-reactivity was limited to heart rate (HR) and electrodermal activity (EDA). EDA indexes sympathetic arousal, which is tightly linked to processing of addiction-related cues^[62,88–90]^. For example, addiction related cues presented in VR increased SCR amplitudes in participants suffering from nicotine addiction ^[89,90]^. In contrast, in the present study there was no significant effect of entering the specific gambling-related or neutral sections of the VR environments (i.e. comparing F5 vs. S1) in either group for any physiological measure. Instead, in both groups, the tonic skin conductance level (SCL) increased substantially upon VR immersion in both groups (effect size *r* between .66 and .81 in both groups and both sessions) and remained elevated until the end of the experiment, an effect we have observed previously in non-gambling controls^[39]^. No corresponding effects were observed for heart rate or the number of spontaneous skin conductance responses. These results suggest a general increase of sympathetic activity during VR immersion, but do not show evidence for physiological cue-reactivity in GD in the present VR setting.

### Behavioral performance

The behavioral tasks replicated earlier results in gamblers and controls^[5,17]^. Regular slot machine gamblers discounted rewards substantially more steeply than controls, and this was the case across model-agnostic (AUC) and model-based measures (i.e. *log(k)* in softmax and diffusion models), resonating with a range of earlier results in behavioral addictions and substance-use disorders ^[1,5,9,10]^. We likewise replicated the recent finding of reduced MB decision-making in participants suffering from GD^[17]^. The model-agnostic analysis of RTs revealed that both groups showed longer S2 RTs after rare transitions, reflecting MB decision-making^[14,17]^. However, this effect was significantly increased in non-gambling controls. These results were also supported by comprehensive computational modelling. A hybrid reinforcement learning (RL) model showed very strong evidence for reduced MB decision making in the gambling group, and this was the case for both softmax and DDM choice rules.

### Behavioral cue-reactivity

In contrast to these robust group differences, and contrasting with our preregistered hypothesis, neither temporal discounting nor model-based RL were substantially modulated by the virtual gambling environment in the GD group. A seminal study by Dixon and colleagues^[27]^ showed increased discounting in gamblers when tested in a gambling environment, an effect that we recently replicated^[28]^. Other studies showed increased temporal discounting^[6]^ or increased risk-taking^[29]^ in participants suffering from GD in the presence of gambling-related visual cues^[6,29,91]^. The group level posterior distribution of the shift parameter *s*_*k*_ of the hyperbolic discounting model indicated only inconclusive evidence for increased discounting in the gambling group, and this was the case for both softmax and DDM choice rules. There was, however, strong evidence for *decreased* discounting in the VR_gambling_ session in the non-gambling control group. There was also strong evidence that this decrease was significantly more pronounced than in gambling group (comparison of the group level posteriors of the *s*_*k*_ parameter). The fact that non-gambling control participants tended to discount rewards less steeply in the VR_gambling_ session is of potential interest, but it is important to note that in a previous study employing the same VR design^[39]^ we observed the reverse effect in a group of non-gambling controls. In that study, controls if anything showed reduced temporal discounting in the VR_neutral_ session. Effects of virtual gambling environment exposure in non-gambling controls are therefore overall inconclusive.

An analysis of the DDM parameters revealed strong evidence for a decrease in boundary separation in the VR_gambling_ session in the GD group. Participants reporting regular slot machine gambling appeared to increasingly trade-off accuracy in favour of speed in the VR_gambling_ session. However, there was only moderate evidence for this decrease being stronger in the gambling group than in the non-gambling control group (comparison of the group level posteriors of the s_*α*_ parameter). It is thus difficult to draw strong conclusion from this observation. Nevertheless, a tendency towards a decrease in boundary separation would indicate that regular slot machine gamblers might attend less to actual value differences between the SS and the LL, instead preferring a more rapid response rate. In addition, the group level posterior distribution of 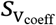 in controls suggests that the value difference between the LL and the SS had a stronger influence in the trial-wise RT in the VR_gambling_ session, indicating that non-gambling control participants placed a stronger weight onto the value differences between SS and LL in the VR_gambling_ session. Again, there was only moderate evidence for this effect being stronger in the non-gambling control group than in the gambling group (comparison of the group level posteriors of the 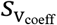 parameter).

Taken together, earlier findings of steeper discounting in gamblers when tested in real gambling venues ^[27]^ which we recently replicated^[28]^, did not translate to VR. Whether this is due to the specific design of the present VR environments, or due to more general effects of VR on cognitive load and/or arousal (see below) is an open question.

We hypothesized that exposure the VR_gambling_ environment would reduce MB and increase of MF decision-making in regular slot machine gamblers. This hypothesis was based on the idea that addiction-related cues might trigger pathological habits^[21,92]^, which in turn could be related to reduced MB decision-making^[16,93]^. An alternative view is that addiction might instead be associated with excessive goal-directed behavior, in particular in the presence of addiction related cues^[94]^. We observed the latter effect in a recent study in regular slot-machine gamblers^[28]^. Regular gamblers showed a substantial increase in model-based control when tested in a gambling venue. However, in the present VR setting, we observed little evidence for either of these effects. Gamblers showed no evidence for longer RTs after rare transitions in the VR_gambling_ session, an index for MB decision-making. The hybrid model parameter *s*_*β*_ showed only moderate evidence for an increase in MB decision-making in the VR_gambling_ environment in both groups. 95% HDIs were overlapping with zero and the change was of similar magnitude in both groups, suggesting the absence of specific cue-reactivity effects on MB decision-making in the gambling group. A similar picture emerged for the corresponding parameters from the hybrid model with DDM choice rule. Taken together, there was no conclusive evidence for VR effects on either MB or MF decision-making in either group. As reductions in MB control constitute potential transdiagnostic markers for compulsivity-related disorders^[16,93]^ it is interesting that the VR_gambling_ environment did not cause effects in the gambling group. Possible explanations for the lack of cue-reactivity effects are discussed below.

### Conclusion

In contrast to the absence of behavioral and physiological cue-reactivity effects, the present VR set-up increased the subjective urge to gamble in participants reporting frequent slot machine gambling. This does not render VR generally unsuitable, as it has been shown here and by other groups that gambling related VR environments can induce craving^[40]^. Rather, behavioral, and physiological cue-reactivity effects might depend on specific VR design features. It is of course possible that different VR designs might have yielded the predicted effects. In addition to the specifics of the VR environments, more general effects of VR immersion might have precluded us from detecting physiological and/or behavioral cue-reactivity effects. In particular, SCL exhibited a substantial overall effect of VR immersion across groups and conditions, replicating our previous observation^[39]^. This might reflect increased cognitive load of VR immersion ^[95,96]^, which could interfere with the expression of behavioral effects of gambling-related environments. Likewise, physiological correlates of VR-related cognitive load could have precluded us from detecting more subtle modulation of SCL due to cue-reactivity effects. The lack of behavioral and/or physiological correlates for the reported increase in subjective craving may warrant caution for future applications of VR in exposure therapy and addiction science. Exposure therapy aims to confront patients with key stimuli and train strategies to overcome craving or fear responses. It is therefore important to note that VR might be limited in ecological validity to reproduce real-life behavioral effects^[27,28]^.

### Limitations

There are several limitations that need to be acknowledged here. First, although groups were matched on key variables, group differences on depressive symptoms and overall psychopathology remained, as in previous studies ^[6,17]^. Second, participants spend between thirty and forty minutes in VR. Behavioral tasks were performed after the initial exploration phase. Participants might therefore have been fatigued or experienced some discomfort during task performance. Future studies might thus benefit from shorter designs. Third, the immersion in VR was constrained by the available physical lab space. The experimenter had to ensure the safety of participant by giving external instructions when needed. Distractions caused by these instructions might have reduced the immersion experienced by the participants. Additionally, such instructions might have affected physiological measures. Future research would benefit from implementing clear markers within the VR environments to ensure safety without breaking immersion. Finally, the virtual slot machines used in our design did not exactly match most recent machines used in local gambling facilities. This might have reduced the level of realism that the VR environment conveyed. However, since an increase in the subjective urge to gamble was observed in the GD group, this indicates sufficient ecological validity to produce subjective craving. Future research could benefit from improved quality of graphical assets, e.g., by creating objects that more closely resemble current video slot machines.

Overall, we reproduced established group differences in decision-making between participants suffering from GD and non-gambling control participants in a VR setting: Participants reporting frequent gambling showed higher levels of temporal discounting and reduced MB decision-making, compared to non-gambling controls. However, we found little evidence for behavioral or physiological effects of virtual gambling environments in the GD group, despite these environments eliciting increased subjective craving in gamblers. Some caution is therefore warranted when applying VR in experimental or therapeutical contexts, as established behavioral effects of gambling environments^[27,28]^ might not generally replicate in VR. Future studies should delineate how cognitive load and ecological validity could be balanced in VR to create a more naturalistic VR experience.

## Acknowledgments

We thank Mohsen Shaverdy and Diego Saldivar for the implementation of the VR environments and task programming and all members of the Peters Lab at the University of Cologne for helpful discussions. This work was supported by Deutsche Forschungsgemeinschaft (DFG, grant PE1627/5-1 to J.P.).

## Supplementary Materials

**Supplementary Table 1.**
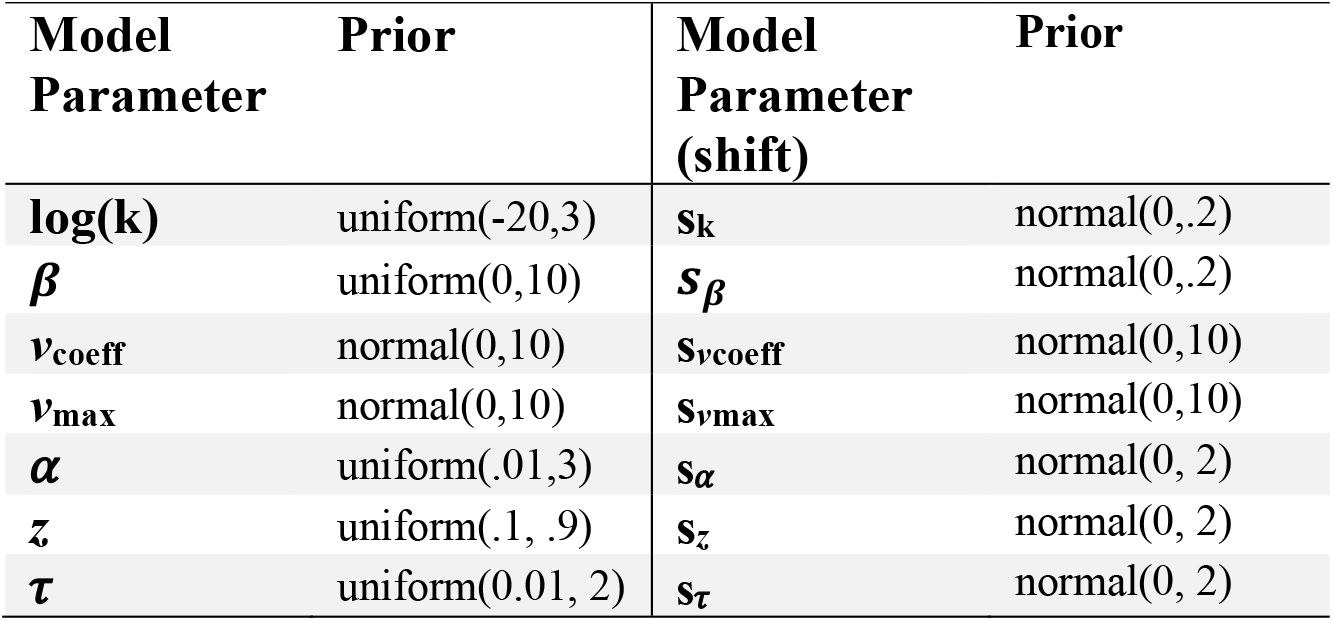
Priors for all parameters used in the computational modelling analysis of the temporal discounting task.

**Supplementary Table 2.** Priors for all parameters used in the computational modelling analysis of the 2-step task.

| Model<br>Parameter | Prior | Model<br>Parameter<br>(shift) | Prior |
| --- | --- | --- | --- |
| $\eta_1$ | uniform(-4,4) | $s\eta_1$ | normal(0, 1) |
| $\eta_2$ | uniform(-4,4) | $s\eta_2$ | normal(0, 1) |
| $\beta_{MB}$ | normal(0,15) | $s\beta_{MB}$ | normal(0,3) |
| $\beta_{MF}$ | normal(0,15) | $s\beta_{MF}$ | normal(0,3) |
| $\beta_2$ | normal(0,15) | $s\beta_2$ | normal(0,3) |
| $p$ | normal(0,10) | $s_p$ | normal(0, 2) |
| $v_{\text{coeff MB}}$ | normal(0,10) | $s_{v_{\text{coeff MB}}}$ | normal(0,10) |
| $v_{\text{coeff MF}}$ | normal(0,10) | $s_{v_{\text{coeff MF}}}$ | normal(0,10) |
| $v_{\text{coeff2}}$ | normal(0,10) | $s_{v_{\text{coeff2}}}$ | normal(0,10) |
| $v_{\text{max1}}$ | normal(0,10) | $s_{v_{\text{max1}}}$ | normal(0,10) |
| $v_{\text{max2}}$ | normal(0,10) | $s_{v_{\text{max2}}}$ | normal(0,10) |
| $\alpha_1$ | uniform(.01,3) | $s\alpha_1$ | normal(0, 2) |
| $\alpha_2$ | uniform(.01,3) | $s\alpha_2$ | normal(0, 2) |
| $\tau_1$ | uniform(.01, 2) | $s\tau_1$ | normal(0, 2) |
| $\tau_2$ | uniform(.01, 2) | $s\tau_2$ | normal(0, 2) |

### Supplemental model-agnostic analysis 2-step task

As described in our preregistration we performed an additional model-agnostic HGLM analysis of the 2-step task. We modelled the probability to repeat the choice from the previous trial (1 if the choice was repeated and 0 if it was not) as a function of the transition in the previous trial (rare or common), the reward received in the previous trial (0 if the reward was lower than the mean of the rewards received in the past 20 trials and 1 otherwise), the group (gambling or non-gambling control), and finally the session (VR_neutral_ vs VR_gambling_).

The results were mostly in line with the results obtained from the other analysis. The effect of Reward (z = -4.024, p < .001) and the interaction Reward*Transition (z = 9.308, p < .001) were significant indicating that overall participants in both groups showed MB and MF decision-making. Additionally, the three-way interaction Trans*Rew*Group was significant (z = -6.389, p < .001) indicating stronger MB decision-making in the non-gambling control group. Finally, none of the terms including the session were significant suggesting the absence of effects specifically caused by the VR environments.

**Supplementary Table 3.**
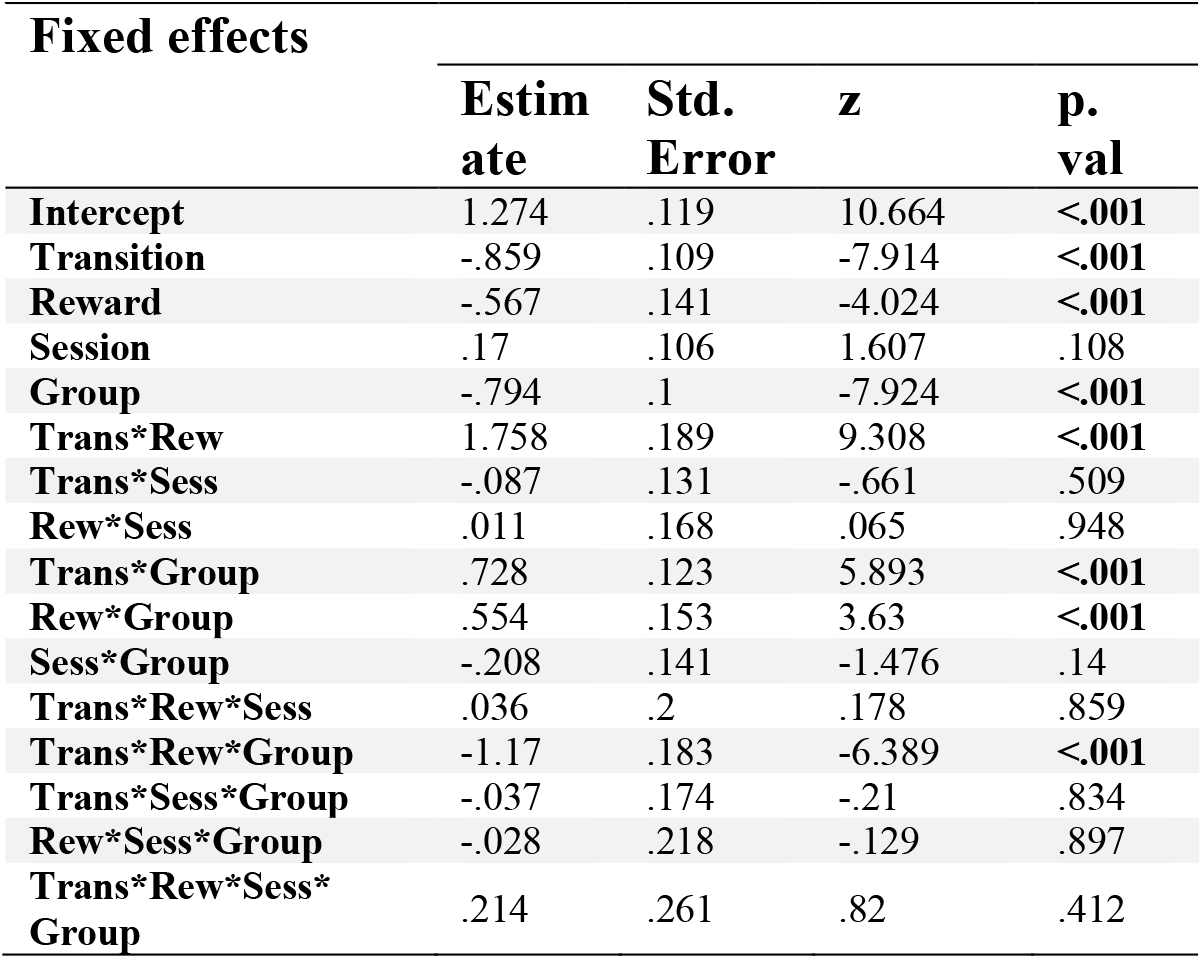
Results of the supplementary HGLM analysis of the 2-step task. The probability to repeat S1 choices (pStay) is modelled as a function of previous transition and reward, the group affiliation and session. p-values printed in bold font are significant at a threshold of .05.

Fixed effects
|  | Estimate | Std. Error | z | p. val |
| --- | --- | --- | --- | --- |
| Intercept | 1.274 | .119 | 10.664 | <.001 |
| Transition | -.859 | .109 | -7.914 | <.001 |
| Reward | -.567 | .141 | -4.024 | <.001 |
| Session | .17 | .106 | 1.607 | .108 |
| Group | -.794 | .1 | -7.924 | <.001 |
| Trans*Rew | 1.758 | .189 | 9.308 | <.001 |
| Trans*Sess | -.087 | .131 | -.661 | .509 |
| Rew*Sess | .011 | .168 | .065 | .948 |
| Trans*Group | .728 | .123 | 5.893 | <.001 |
| Rew*Group | .554 | .153 | 3.63 | <.001 |
| Sess*Group | -.208 | .141 | -1.476 | .14 |
| Trans*Rew*Sess | .036 | .2 | .178 | .859 |
| Trans*Rew*Group | -1.17 | .183 | -6.389 | <.001 |
| Trans*Sess*Group | -.037 | .174 | -.21 | .834 |
| Rew*Sess*Group | -.028 | .218 | -.129 | .897 |
| Trans*Rew*Sess*Group | .214 | .261 | .82 | .412 |

**Supplementary Table 4.** Results of Wilcoxon signed rank tests of the SCL data for the gambling group. p-values printed in bold font are significant at a Bonferroni corrected threshold of .004.

Comparison
|  | Z | df | p. val | r |
| --- | --- | --- | --- | --- |
| B5 vs F1<br>VR<sub>neutral</sub> | -4.099 | 30 | <.001 | .804 |
| B5 vs F1<br>VR<sub>gambling</sub> | -3.552 | 30 | <.001 | .661 |
| VR<sub>neutral</sub> –<br>VR<sub>gambling</sub> | -.683 | 30 | .495 | -.134 |
| F5 vs S1<br>VR<sub>neutral</sub> | -2.141 | 30 | .032 | -.42 |
| F5 vs S1<br>VR<sub>gambling</sub> | .729 | 30 | .466 | .143 |
| VR<sub>neutral</sub> –<br>VR<sub>gambling</sub> | -1.753 | 30 | .008 | -.344 |

**Supplementary Table 5.** Results of Wilcoxon signed rank tests of the SCL data for the control group. p-values printed in bold font are significant at a Bonferroni corrected threshold of .004.

| Comparison | Z | df | p. val | r |
| --- | --- | --- | --- | --- |
| B5 vs F1<br>VR<sub>neutral</sub> | -3.552 | 28 | **<.001** | .697 |
| B5 vs F1<br>VR<sub>gambling</sub> | -3.917 | 28 | **<.001** | .768 |
| VR<sub>neutral</sub> –<br>VR<sub>gambling</sub> | 1.139 | 28 | .255 | .223 |
| F5 vs S1<br>VR<sub>neutral</sub> | -1.389 | 28 | .165 | -.272 |
| F5 vs S1<br>VR<sub>gambling</sub> | 1.389 | 28 | .165 | .272 |
| VR<sub>neutral</sub> –<br>VR<sub>gambling</sub> | -1.731 | 28 | .084 | -.339 |

**Supplementary Table 6.** Results of Wilcoxon signed rank tests of the nSCRs data for the gambling group. p-values printed in bold font are significant at a Bonferroni corrected threshold of .004.

| Comparison | Z | df | p. val | r |
| --- | --- | --- | --- | --- |
| B5 vs F1<br>VR<sub>neutral</sub> | -.25 | 30 | .802 | -.049 |
| B5 vs F1<br>VR<sub>gambling</sub> | -2.811 | 30 | .005 | -.551 |
| VR<sub>neutral</sub> –<br>VR<sub>gambling</sub> | 1.684 | 30 | .092 | -.33 |
| F5 vs S1<br>VR<sub>neutral</sub> | -.713 | 30 | .476 | -.139 |
| F5 vs S1<br>VR<sub>gambling</sub> | 1.116 | 30 | .265 | .219 |
| VR<sub>neutral</sub> –<br>VR<sub>gambling</sub> | -1.048 | 30 | .295 | -.205 |

**Supplementary Table 7.** Results of Wilcoxon signed rank tests of the SCL data for the control group. p-values printed in bold font are significant at a Bonferroni corrected threshold of .004.

| Comparison | Z | df | p. val | r |
| --- | --- | --- | --- | --- |
| B5 vs F1<br>VR<sub>neutral</sub> | .888 | 28 | .375 | .174 |
| B5 vs F1<br>VR<sub>gambling</sub> | .114 | 28 | .891 | .027 |
| VR<sub>neutral</sub> –<br>VR<sub>gambling</sub> | .137 | 28 | .89 | .025 |
| F5 vs S1<br>VR<sub>neutral</sub> | -1.776 | 28 | .076 | -.348 |
| F5 vs S1<br>VR<sub>gambling</sub> | -2.852 | 28 | .004 | -.559 |
| VR<sub>neutral</sub> –<br>VR<sub>gambling</sub> | .433 | 28 | .665 | .085 |

**Supplementary Table 8.** Results of Wilcoxon signed rank tests of the HR data for the gambling group. p-values printed in bold font are significant at a Bonferroni corrected threshold of .004.

| Comparison | Z | Df | p. val | r |
| --- | --- | --- | --- | --- |
| B5 vs F1<br>VR<sub>neutral</sub> | -2.664 | 30 | .008 | -.523 |
| B5 vs F1<br>VR<sub>gambling</sub> | -3.53 | 30 | **< .001** | -.692 |
| VR<sub>neutral</sub> –<br>VR<sub>gambling</sub> | -.387 | 30 | .699 | -.076 |
| F5 vs S1<br>VR<sub>neutral</sub> | -1.001 | 30 | .316 | -.197 |
| F5 vs S1<br>VR<sub>gambling</sub> | .182 | 30 | .855 | -.036 |
| VR<sub>neutral</sub> –<br>VR<sub>gambling</sub> | -.729 | 30 | .4662 | -.143 |

**Supplementary Table 9.** Results of Wilcoxon signed rank tests of the HR data for the control group. p-values printed in bold font are significant at a Bonferroni corrected threshold of .004.

| Comparison | Z | df | p. val | r |
| --- | --- | --- | --- | --- |
| B5 vs F1<br>VR<sub>neutral</sub> | .615 | 28 | .539 | .121 |
| B5 vs F1<br>VR<sub>gambling</sub> | -1.389 | 28 | .165 | -.272 |
| VR<sub>neutral</sub> –<br>VR<sub>gambling</sub> | 1.23 | 28 | .219 | -.241 |
| F5 vs S1<br>VR<sub>neutral</sub> | -.046 | 28 | .964 | -.009 |
| F5 vs S1<br>VR<sub>gambling</sub> | 1.526 | 28 | .127 | .299 |
| VR<sub>neutral</sub> –<br>VR<sub>gambling</sub> | -.66 | 28 | .509 | -.13 |

